# Learning the dynamic organization of a replicating bacterial chromosome from time-course Hi-C data

**DOI:** 10.1101/2025.04.03.647011

**Authors:** Janni Harju, Joris J.B. Messelink, Lucas Tröger, Grzegorz Gradziuk, Imesha Rathnayaka, Maria Billini, Martin Thanbichler, Muriel C.F. van Teeseling, Chase P. Broedersz

## Abstract

Bacterial chromosomes are in continual motion as they undergo concurrent transcription, replication, and segregation. Time-course Hi-C experiments hold promise for studying chromosome organization across the cell cycle, but interpreting Hi-C data from dynamic systems remains challenging. Here, we develop a fully data-driven 4D Maximum Entropy approach to extract a model for the dynamic organization of a replicating bacterial chromosome directly from time-course Hi-C and microscopy data. After validating our 4D data-driven model for *Caulobacter crescentus* against independent microscopy data, we infer quantitative information about changes in chromosome organization across the bacterial replication cycle. Our model reveals a sustained global linear organization of the *C. crescentus* chromosome during replication, as well as dynamic patterns of local chromosome extension induced by the replication forks. We use these data-driven inferences to constrain a mechanistic model for a replicating bacterial chromosome. Our model demonstrates that origin-pulling by a ParAB*S* -like system, together with loop extrusion by condensin, can explain our inferred large-scale chromosome segregation patterns. The inferred replication-induced local changes in chromosome compaction, however, require additional mechanisms, which we attribute to replication-induced NAP unbinding and positive supercoiling. Overall, our work introduces a rigorous data-driven framework for quantitatively interpreting time-course Hi-C data, and offers new mechanistic insights into bacterial chromosome organization across the cell cycle.

---

Bacterial chromosome organization is intrinsically dynamic, with processes such as transcription, replication, and segregation often coinciding for most of the cell cycle [1–3]. Open questions remain about the dynamic interplay between these processes and different chromosome organization mechanisms, including the abundant Nucleoid Associated Proteins (NAPs) [4, 5], and the highly conserved Structural Maintenance of Chromosome (SMC) complexes, which fold the chromosome via an active loop extrusion process [6–12]. However, studying bacterial chromosome organization at any replication stage, let alone throughout the cell cycle, remains challenging: since bacterial chromosomes are typically compressed by orders of magnitude to fit into micron-sized cells, microscopy studies offer limited spatial resolution, and can typically only track a few labeled loci simultaneously. Alternatively, chromosome organization can be probed using chromosome conformation capture methods, such as Hi-C, yielding population-averaged counts of how often pairs of chromosomal loci are spatially proximate [13, 14]. However, although Hi-C studies have increased our understanding of chromosome organization in many bacteria [6, 15–18], it remains challenging to use these data to study bacterial chromosome dynamics.

A major challenge in studying chromosome organization is linking population-averaged Hi-C counts to the underlying distribution of 3D chromosome configurations [19, 20]. This problem is further exacerbated when Hi-C data are gathered from asynchronous bacterial populations, where contact counts are averaged over cells at different replication stages. Previous studies introduced the first data-driven approaches for modeling replicating bacterial chromosomes, using asynchronous Hi-C data to construct models for several discrete replication stages [21, 22]. However, it remains untested whether asynchronous Hi-C data contain enough information to infer how chromosome organization changes over time. More recently, time-course Hi-C data were used to infer a model for mitotic chromosome organization in chicken cells [23], but this model did not account for chromosome replication or segregation. Importantly, none of these previous approaches took into account a crucial feature of replicating systems: Hi-C scores are typically averaged over contacts between locus pairs on the same dsDNA polymer strand (*cis*) and on different replicated strands (*trans*). Neglecting *trans*-contacts is a major assumption, which has not been validated experimentally.

Thus, the field still lacks a principled method to construct a minimal and unbiased data-driven model for replicating chromosomes that would faithfully reproduce input Hi-C data.

Here, we introduce a data-driven 4D Maximum Entropy (4D-MaxEnt) approach that uses information theory and time-course Hi-C data on synchronized cell populations to resolve the changing ‘landscape’ of chromosome organization in replicating bacteria. We constrain our model on time-course Hi-C data [14] and microscopy data for the origins of replication from replicating *Caulobacter crescentus* cells (Fig. 1A). Like many bacteria, *C. crescentus* has a single, circular chromosome. When replication begins, two replication forks start moving in opposite directions from the origin of replication (*ori*) towards the terminus (*ter*) [24] (Fig. 1D). Throughout this process, one *ori* remains tethered to a cell pole [25, 26], whereas the other is actively transported by the ParAB*S* system towards the opposite pole [27, 28].

**FIG. 1.**
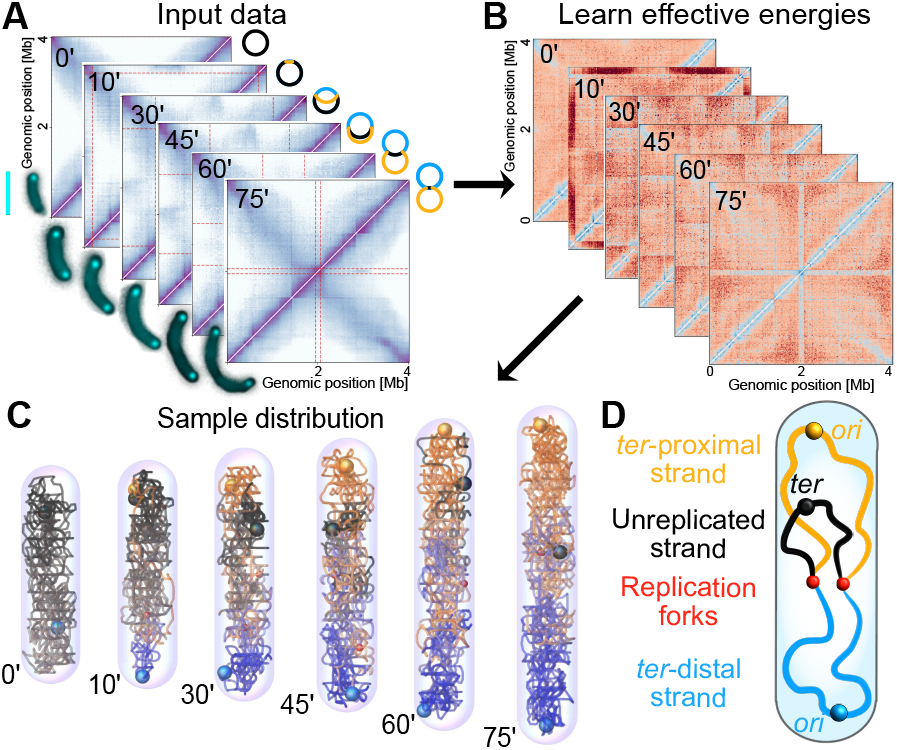
Description of the 4D-MaxEnt model. (A) We constrain our model on time-course Hi-C data from [14] (SI Appendix, Fig. S1) as well as the mean separation between replicated origins, extracted from microscopy images (scale-bar 2 µm). Shown are exemplary cells from strain Tn4 with a labeled *oriC* in cyan (SI Appendix, Table S1). (B) We infer the effective energies of the model using gradient descent until the model contact map has a *>* 98% Pearson correlation with the input contact map, as seen between replicate Hi-C experiments [14]. (C) Given the converged effective energies, we sample the chromosome configuration distribution using Monte Carlo simulations. (D) Cartoon of a replicating *C. crescentus* chromosome. The replicated strand with its *ori* closer to the *ter* is defined as “*ter* -proximal”.

Our 4D-MaxEnt model enables us to sample single-cell 3D chromosome configurations across the cell cycle (Fig. 1C). After validating the model against independent microscopy data, we make quantitative inferences about large-scale chromosome organization throughout the replication cycle. The 4D-MaxEnt model shows early displacement of unreplicated regions to mid-cell, as well as sustained linear organization of the *C. crescentus* chromosome, consistent with previous experimental reports [29, 30]. However, we also find subtle but significant deviations from linear organization at intermediate replication stages, indicating uneven compaction of the chromosome. To explain these findings, we use our data-driven inferences to constrain a simple mechanistic model for replicating bacterial chromosomes. We find that a ParAB*S* -like origin-pulling mechanism is sufficient to segregate newly replicated regions and to establish linear order of the chromosome, consistent with the null hypothesis that ParAB*S* is the main driver of chromosome segregation in *C. crescentus*. Loop-extruders, however, contribute to deviations from linear chromosome organization and the early displacement of unreplicated chromosomal regions in our mechanistic model. Owing to its high spatial resolution, the 4D-MaxEnt model reveals a pattern of local extension behind and compaction ahead of the replication forks, consistent with NAP unbinding [31] and supercoiling changes [32, 33] during replication. Our work illustrates the wealth of quantitative information that can be inferred from time-course Hi-C data and how mechanistic modeling can help interpret data-driven inferences.

## MODEL

Our goal is to use time-course Hi-C data to infer a probability distribution for the 3D configurations of a bacterial chromosome at different replication stages. While Hi-C data measure population-averaged contact frequencies, which can be challenging to interpret directly, a distribution of 3D chromosome configurations can be used to infer interpretable 3D conformational statistics over the cell cycle.

To infer a statistical model for chromosome conformations, we must solve a uniqueness issue inherent to Hi-C-based inference: many distributions can be consistent with the same Hi-C data, so how should we select a model in an unbiased way? We select the minimal least-constrained model by employing an information theoretical approach based on the Maximum Entropy (MaxEnt) principle [34]. The Shannon entropy *S*[*P*(*x*)] of a distribution *P*(*x*) measures the average uncertainty of sampling outcomes, reflecting the lack of structure in the distribution. To illustrate, for any given set 𝒮, a uniform distribution over𝒮 has maximal entropy, whereas a highly peaked distribution has low entropy. A MaxEnt model is constructed by maximizing *S*[*P*(*x*)] under experimental constraints, e.g. Hi-C data [35–38]. Thus, the MaxEnt principle allows us to select the unique, maximally uncertain distribution explaining the experimental data.

In our previous work [38], we introduced a MaxEnt approach for modeling the organization of an unreplicating bacterial chromosome based on Hi-C data. Here, we construct a 4D-MaxEnt model for growing, replicating bacteria, taking into account: 1) the topology of the replicating chromosome; 2) cell growth; and 3) the experimental Hi-C constraints at different times, which represent relative contact counts averaged over *cis*- and *trans*-locus pairs.

### Defining the time-dependent phase space

To construct the 4D-MaxEnt model for replicating bacterial chromosomes, we first define the model distribution’s time-dependent domain or “phase space” 𝒮 (*t*): the set of all possible 3D chromosome configurations at a time-point *t*. At a given time-point, the possible states of the system are the 3D configurations of a partially replicated chromosome, confined to a growing cell. To define this phase space mathematically, we use a coarse-grained representation of the chromosome as a lattice polymer of *N* monomers, confined to a cylinder. The coarse-graining scale is set by the resolution of the Hi-C experiments: each 10 kb Hi-C bin corresponds to a segment of the polymer. To calibrate the spatial scale of the model, we previously used microscopy experiments to determine the distance distribution for loci separated by 10 kb [38]. We then determined that a monomer length (or lattice spacing) of 88 nm gives good agreement between the distance distributions of the model and experiment. Since we can resolve the spatial separation between two loci in our model when they are at least a lattice spacing apart, the monomer length also defines the spatial resolution of our model.

The available states of the system depend on the replication stage. At time *t* from the start of replication, we include two replication forks at genomic positions *F*_1_(*t*) *< F*_2_(*t*), relative to the origin at position 1, and attach a replicated segment of genomic length *F*_1_ + *N* −*F*_2_ between these sites. Following [14], the mean positions of the replication forks at each time-point are inferred directly from the Hi-C data (SI Appendix, Fig. S2). Once the fork positions for all time-points are determined, a chromosome configuration {**r, r**^*′*^} is defined by the lattice positions of all unreplicated monomers (**r**_*i*_; *F*_1_ *< i < F*_2_),and all pairs of replicated monomers (**r**_*j*_,**r**^*′*^_*j*_; *j* ≤ *F*_1_ or *j*≥ *F*_2_). This partially replicated chromosome is confined to a cylinder with rounded caps, with the confinement dimensions at each time-point set using microscopy data (SI Appendix, Fig. S3A).

### Imposing Hi-C constraints

Given our time-dependent phase-space 𝒮 (*t*) ∋{**r, r**^*′*^}, we will now define a time-dependent 4D-MaxEnt distribution *P*_ME_ ({**r, r**^*′*^}, *t*). In practice, at each time-point *t*, we seek a probability distribution for 3D chromosome configurations consistent with Hi-C data. This distribution is derived by maximizing the Shannon entropy under the constraint that the model’s pairwise contact frequencies, averaged over *cis*- and *trans*-contacts, are proportional to the bias-corrected Hi-C counts (SI Appendix):

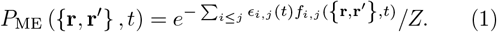

Here *Z* is the partition function, *ϵ*_*i,j*_ is the effective contact energy for monomers *i* and *j*, and *f*_*i,j*_ is a function that counts the total number of *cis*- and *trans*-contacts between genomic regions *i, j*:

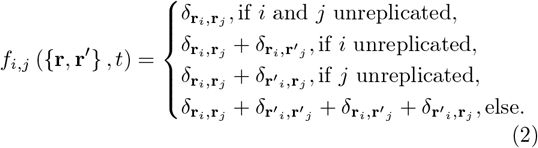

The Kronecker delta-function *δ*_**r***i*,**r***j*_ is equal to 1 if monomers *i* and *j* occupy the same lattice site, and zero otherwise. In other words, there is an effective energy associated with monomers *i* and *j* (or their replicates) co-localizing. At each time point, our model is mathematically equivalent to an equilibrium polymer with close-range pairwise interactions between monomers.

Since Hi-C experiments yield relative contact counts rather than absolute contact frequencies, there is an unknown constant of proportionality *C*(*t*) between the model’s contact frequencies and the measured Hi-C counts. We set this constant in a data-driven way by maximizing the distribution entropy relative to *C*(*t*) (SI Appendix). Our approach thus does not rely on assuming a constant of proportionality between Hi-C counts and contact probabilities [22, 35, 36], on converting Hi-C counts to mean pairwise distances between loci [19, 21, 39, 40], or making assumptions about *trans*-interactions between chromosomes [21, 22].

Given (1), the remaining task is to infer the effective energies *ϵ*_*i,j*_(*t*) from time-course Hi-C data. We achieve this using a gradient descent algorithm (SI Appendix), where the effective energies are iteratively adjusted until the model matches input data [38]. We then sample the model ensemble using Monte Carlo simulations [38], without assuming that the chromosome evolves at thermal equilibrium. This contrasts previous works where data-driven potentials were incorporated into molecular dynamics simulations to infer chromosome configuration time-trajectories [22, 35–37]. Furthermore, since each Hi-C map in the time-course is mapped to a separate set of effective energies *ϵ*_*i,j*_(*t*) (SI Appendix, Fig. S4A), our approach allows the ‘landscape’ of chromosome organization to change over time.

### Hi-C and *ori-ori* distance constraints are sufficient to constrain the model

When constructing a MaxEnt model, it is *a priori* unclear whether a given set of constraints will give rise to a model with enough predictive power. We hence start by checking whether Hi-C constraints alone are sufficient to constrain a predictive model for bacterial chromosome organization during replication.

We find that a model only constrained on Hi-C data shows replicated chromosomal strands co-localizing rather than segregating (SI Appendix, Fig. S3C,D).By contrast, experiments have shown that *C. crescentus* chromosomes are simultaneously replicated and segregated [30]. Sister co-localization in this model is caused by Hi-C experiments not discriminating between *cis*- and *trans*-contacts, which can lead to attraction between chromosome replicates in the inferred model (SI Appendix, Fig. S4A). We conclude that Hi-C data alone do not contain sufficient information to infer a model for segregating bacterial chromosomes.

To construct an informative model for replicating chromosomes, we must introduce an additional constraint that, at least locally, discriminates between *cis*- and *trans*-contacts. Motivated by how the ParAB*S* system initiates *C. crescentus* chromosome segregation by pulling the origins of replication to opposite cell poles [27, 28], we constrain the mean long-axis separation between replicated *ori* regions to match experimental data (SI Appendix). In this case, the 4D-MaxEnt model infers chromosome segregation at all replication stages. Hence, by introducing one additional constraint on the *ori-ori* distance, we construct our 4D-MaxEnt model that is both consistent with experimental Hi-C data and with experimental evidence for concurrent chromosome replication and segregation in *C. crescentus*.

## RESULTS

### The 4D-MaxEnt model is validated by imaging data

Having obtained the least-assuming data-driven model for replicating bacterial chromosomes, we can now sample single-cell chromosome configurations from the resulting 4D-MaxEnt distribution to infer quantitative chromosome organization statistics throughout the bacterial cell cycle.

To validate our 4D-MaxEnt model, we infer the mean long-axis positions of several loci over time, and compare with independent microscopy data that were not used to constrain the model (Materials and Methods, Fig. 2A). This allows us to test whether the model can accurately capture 4D chromosome organization. Specifically, we use our model to calculate the distance from each unreplicated locus to its nearest cell pole, and for each pair of replicated loci, we calculate their distances to the closest pole to either locus (Fig. 2C). In experiments, the microscope’s resolution introduces a systematic bias in inter-locus distance measurements, as short distances between two replicated loci cannot be resolved (SI Appendix, Fig. S5). Thus, to fairly compare the model with experiments, we calculate bias-corrected 4D-MaxEnt distances between loci (SI Appendix).

**FIG. 2.**
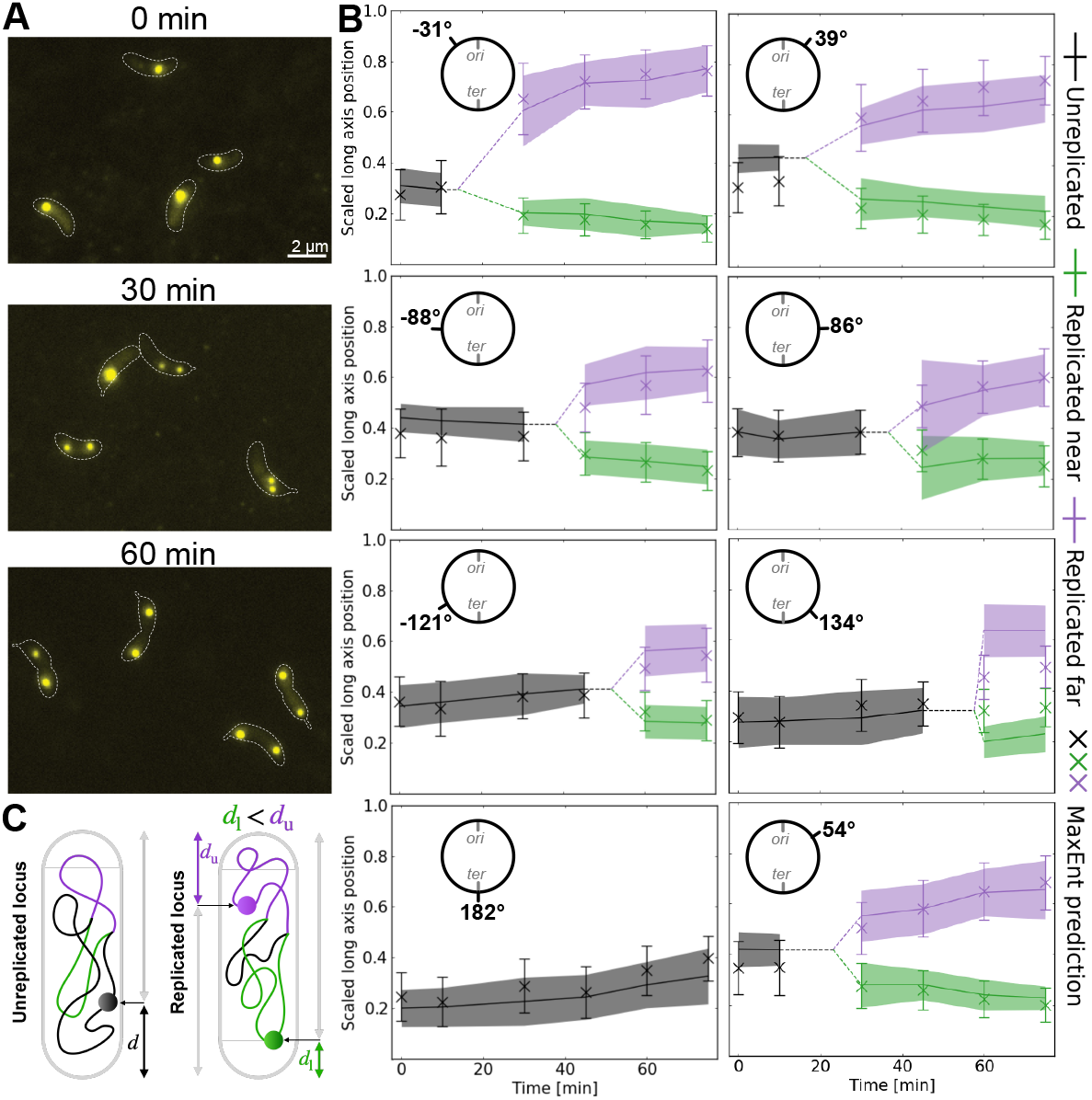
Validation of the model. (A) Representative microscopy images of cells of strain Tn102 (SI Appendix, Table S1) at indicated times after synchronization displaying fluorescently labeled chromosomal loci at -31^°^ in yellow. White dashed lines indicate cell borders. (B) Mean long-axis positions of loci inferred using the 4D-MaxEnt model (crosses, errors bars indicate std.), along with experimental measurements (lines, colored border indicates std.). Dashed lines indicate the predicted time of replication based on a linear replication speed. The 4D-MaxEnt inferences have been corrected for the bias due to the resolution of the experiments (SI Appendix). (C) For all unreplicated loci, we consider their distance to the nearest pole. For replicated loci, we define *d*_*l*_ as the minimum distance between either locus and either pole, and measure the distance to the other locus from this pole.

The bias-corrected 4D-MaxEnt correctly predicts both the means and standard deviations of locus positions across the replication cycle (Fig. 2B; SI Appendix, Fig. S6). This implies that our model captures not only average chromosome conformations, but also variability between individual cells. The 4D-MaxEnt model only slightly underestimates the inter-locus distance after replication for one tagged locus, at 134^°^(third row, second column, Fig. 2B). This region also stands out in the experimental data: replicated loci at 134^°^ segregate extremely rapidly, much faster than the loci at -121^°^ on the opposite chromosomal arm. To understand to what extent our inferred localization profiles are a consequence of *ori* separation constraints, we also infer a model constrained only on the *ori* separations, but not Hi-C data. Since the full model outperforms the *ori* - constrained model in predicting validation data (SI Appendix, Fig. S7), we conclude that our inference procedure successfully translates time-course Hi-C data into interpretable information about the spatial and temporal organization of chromosomes in single cells.

### Inferences reveal segregation patterns and deviations from global linear order

Next, we use our validated 4D-MaxEnt model to gain insight into the global long-axis organization of the *C. crescentus* chromosome across the cell cycle. The inferred mean long-axis positions of all loci reveal the establishment of linear order on newly replicated chromosomal regions (Fig. 3A). The two replicated strands (blue and orange) stretch out in opposite directions from the replication forks, consistent with previous experiments in slower growth conditions [30]. This ensures that, before cell division, each sister chromosome establishes *ori-ter* organization, with its *ori* at a cell pole and the *ter* mid-cell. Importantly, we note that the 4D-MaxEnt model’s mean long-axis position curves show curvature near the replication forks and the cell poles (Fig. 3A), indicating previously unreported deviations from linear organization during replication. Our model also provides further insights into the segregation dynamics of terminal regions. We find that unreplicated regions (black) maintain nearly linear organization across the cell length, as well as their long-axis position relative to mid-cell, until replication completion.

**FIG. 3.**
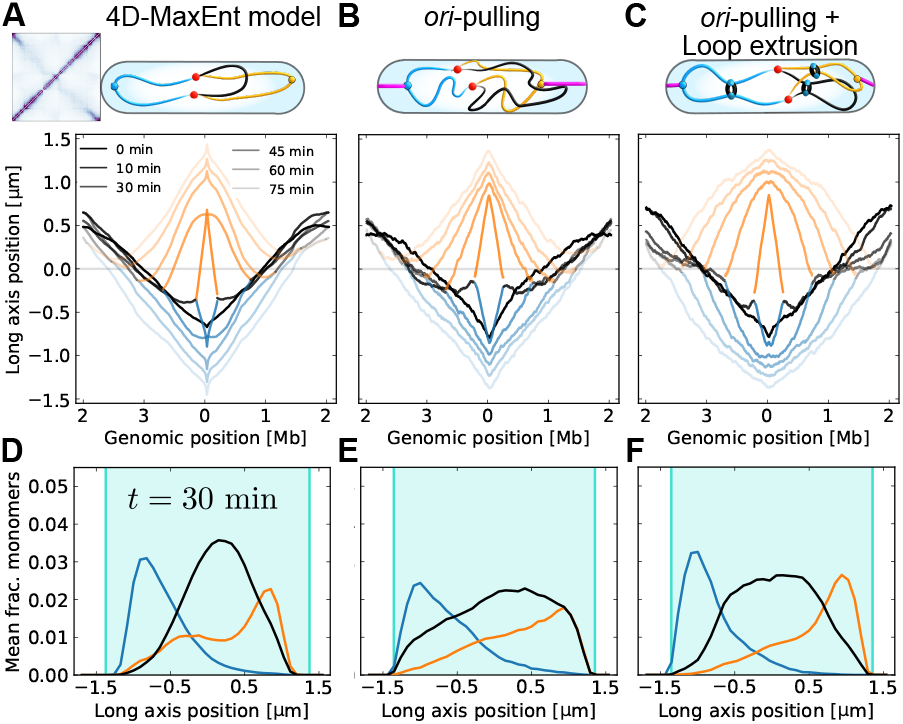
4D-MaxEnt model infers persistent linear chromosome organization and segregation of terminal regions to mid-cell. (A) Inferred mean long axis positions of loci from the 4D-MaxEnt model. *ori*-distal and *ori* - proximal replicated strands in blue and orange, unreplicated strand in black (Fig. 1D). The gray line for zero is defined as mid-cell for all replication stages. At least 4000 samples were used for each time-point. (B) Mean long axis positions of loci from mechanistic model with *ori*-pulling. At least 200 samples were used for each time-point. (C) Mean long axis positions of loci from mechanistic model with both *ori*-pulling and 40 loop-extruders per chromosome, loaded primarily at the origins of replication. (D) Histograms showing relative density of each strand across the length of the confinement in the 4D-MaxEnt model, at *t* = 30 min. Turquoise area indicates confinement length. (E) Density histograms from mechanistic model with *ori*-pulling. (F) Density histograms from mechanistic model with both *ori*-pulling and 40 loop-extruders per chromosome with processivity *λ* = 1200 kb.

To further study the inferred segregation patterns, we construct histograms for the relative densities of the three chromosomal strands along the long axis of the confinement (Materials and Methods). From *t* = 30 min, the 4D-MaxEnt model shows good segregation of the replicated strands, which mainly occupy opposite cell poles (Fig. 3D, SI Appendix, Fig. S8A). The unreplicated region has the highest probability density close to mid-cell, between the two replicated strands. This inference is consistent with the *ter* region moving to mid-cell within 20 minutes of replication initiation [29]. We hence find that the 4D-MaxEnt procedure allows us to make quantitative inferences about chromosome segregation patterns directly from Hi-C data.

### Constraining mechanistic models using Hi-C based inferences

Which biological mechanism(s) could explain the sustained linear organization and density profiles inferred using the 4D-MaxEnt model? In addition to the ParAB*S* system, known to be critical for segregation and the establishment of *ori-ter* order [28], the loop-extruding SMC complex condensin also plays a role in *C. crescentus* chromosome organization and segregation [8]. Condensin attaches onto DNA at a single point, and then actively reels in a loop [7, 9]. In *C. crescentus* and many other bacterial species, condensins are preferentially loaded at the origin of replication, and consequently tie together or “zip-up” the two chromosomal arms as they proceed from *ori* to *ter* [7, 8]. *To test whether the ParABS* system alone could give rise to the 4D-MaxEnt chromosome organization and segregation patterns, or whether loop-extruders are also necessary, we use our data-driven in-ferences to constrain a mechanistic model. Specifically,we perform Molecular Dynamics (MD) simulations of a replicating bacterial chromosome with ParAB*S* -like *ori* - pulling forces and/or loop-extruders [11, 41, 42] (Materials and Methods).

First, we only include *ori*-pulling without loop-extruders in our mechanistic model, corresponding to the null hypothesis that the ParAB*S* system is the main driver of *C. crescentus* chromosome organization. We find that both replicated strands of the chromosome show linear organization, as inferred using the 4D-MaxEnt method (Fig. 3B). Such linear order is not seen in mechanistic simulations without origin-pulling (SI Appendix, Fig. S8, S9), consistent with the ParAB*S* system being critical for large-scale chromosome order. However, the mechanistic model with only *ori*-pulling also shows some deviations from our data-driven inferences: the mean long-axis position profiles do not show curvature near the origins, indicating that the *ori*-proximal regions are more stretched out along the cell’s long axis than in the 4D-MaxEnt model (Fig. 3A). In addition, the density histograms of the model with only *ori*-pulling do not show clear localization of the unreplicated strand (Fig. 3E, SI Appendix, Fig. S8C), in contrast with the 4D-MaxEnt model. These deviations suggest that additional mechanisms are necessary to explain the large-scale organization and segregation patterns inferred using the 4D-MaxEnt model.

Next, we include both *ori*-pulling and loop-extruders in our mechanistic model. Now, the model predicts both linear organization of the replicated strands, as well as curvature of the mean long axis position curves near the *ori* s (Fig. 3C). Intuitively, loop-extruder mediated compaction is expected to bring distant genomic regions closer to the tethered origins, and thus decrease the slope of the mean-long axis curve. Since more compacted regions should show shallower slopes, relatively higher compaction of the origin regions could explain the curvature of the mean long axis curves. We additionally find that a mechanistic model with both origin-pulling and loop extrusion predicts unreplicated regions localizing mid-cell from *t* = 30 min (Fig. 3F, SI Appendix, Fig. S8D), as in the 4D-MaxEnt inferences. These results are consistent with the segregation model we proposed in our previous work, which showed that although origin-pulling can efficiently segregate bacterial chromosomes at early replication stages, additional mechanisms can be needed to segregate terminal regions [11]. We hence conclude that origin-pulling and loop extrusion are sufficient mechanisms to explain the large-scale organization and segregation patterns inferred using the 4D-MaxEnt model.

### Replication forks induce dynamic chromosome compaction and extension

We now turn to smaller scales, and use the 4D-MaxEnt model to study the local extension of genomic regions during replication at a 40-kb scale, limited only by the Hi-C experiment’s resolution. This allows us to infer information about structural modifications of bacterial chromosomes during replication, at a length-scale that is difficult to resolve using microscopy.

As a measure of the local extension of a chromosome at the *n* monomer-scale, we determine

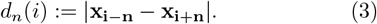

Unless otherwise stated, we consider *n* = 8, corresponding to 20 kb, but our findings are qualitatively similar for values between 10 and 50 kb (SI Appendix, Fig. S10A). To study changes in chromosome compaction during replication, we calculate the relative extension compared to *t* = 0, given by

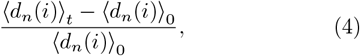

where ⟨… ⟩_*t*_ indicates a mean calculated at time *t*.

At *t* = 10 minutes, the 4D-MaxEnt model shows regions on the newly replicated *ter* -proximal strand extending by up to 60% (SI Appendix, Fig. S10B), indicating that replication and segregation can cause large deformations of the chromosome. Intriguingly, we also find that the 4D-MaxEnt inferences exhibit a persistent trend of chromosome compression ahead of, and extension behind the replication forks (Figure 4A), by factors of roughly 0.95 and 1.1 respectively. Since the data-driven *ori*-constrained model without Hi-C constraints does not show similar compaction changes (SI Appendix, Fig. S11A), we conclude that the 4D-MaxEnt model infers these extension patterns directly from Hi-C data (Materials and Methods).

**FIG. 4.**
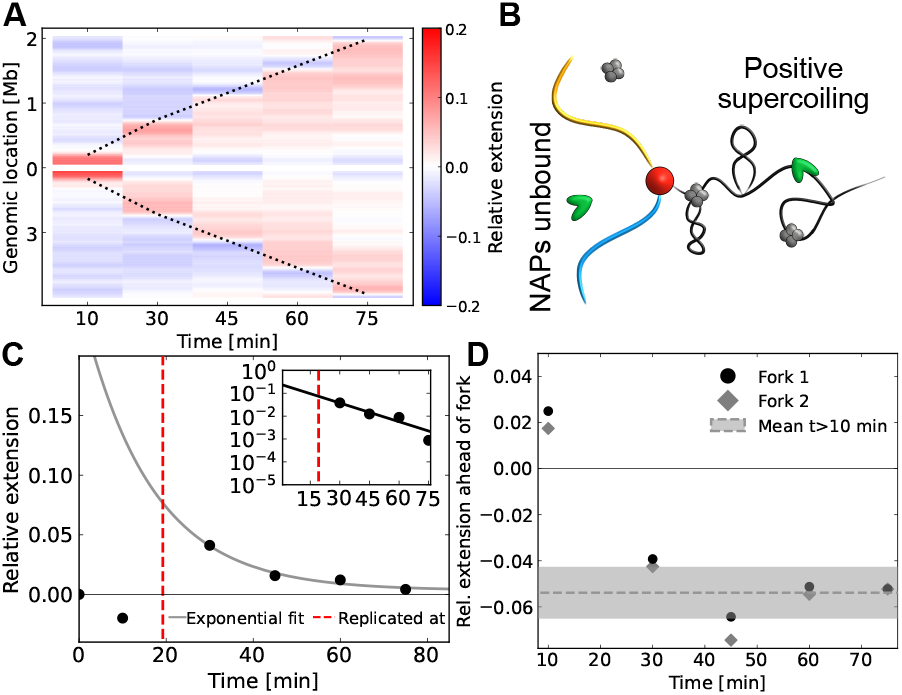
4D-MaxEnt model infers local compression ahead and local extension behind replication fork. (A) Kymograph of relative extension levels inferred using the 4D-MaxEnt model, with *n* = 20 kb. Black dashed lines indicate replication fork positions. (B) Cartoon of potential causes of extension and compaction around replication forks. Replication is expected to cause positive supercoiling ahead of the replication forks, and NAP unbinding behind the forks. (C) Mean relative extension as a function of time for a 200 kb region around a locus at 36°. Red dashed line indicates approximate time of replication. Gray line shows an exponential fit to time-points after replication. The fit time-scale is 16 minutes. (D) Mean relative extension as a function of time in a 100 kb region ahead of the replication forks. Gray line shows mean and standard deviation of data-points for *t ≥* 10.

After replication, the relative extensions averaged over 200 kb regions relax with an exponential decay over time (Fig. 4C, SI Appendix, Fig. S12A), with a mean rate of 1*/*11 min^*−*1^. Varying the averaging length-scale yields qualitatively similar results, with mean decay times between 10-20 minutes (SI Appendix, Fig. S12A). This time-scale implies a genomic length ∼300 kb (replication fork speed times the decay time) across which extension is visible in the wake of the replication fork. The inferred extension behind the replication forks can hence be understood as a local perturbation of chromosome compaction during replication, followed by equilibration on a time-scale of roughly 10-20 minutes.

Similarly, we can study the inferred compaction pattern by averaging the relative compaction over a ∼50 − 100 kb region ahead of the replication forks. We find that this region is compacted for all time-points *t >* 10 minutes (Fig. 4D, SI Appendix, Fig. S12B). The levels of compaction ahead of the forks are similar on both arms of the chromosome, and plateau after ∼10 minutes (Fig. 4D).

Finally, we use our mechanistic model to test whether *ori*-pulling and/or loop extrusion could mechanistically explain the inferred patterns of local extension around the replication forks. We find that *ori*-pulling causes some extension near the origins of replication at *t* = 10 minutes (SI Appendix, Fig. S11A). Loop-extruders, by contrast, tend to slightly compress the chromosome due to an increased density of loops of size *<* 2*n* during replication (SI Appendix, Fig. S11B-E). Importantly, however, neither mechanism results in extension behind and compression ahead of the replication forks, and we hence argue that these inferred patterns should be explained by other biophysical mechanisms.

### NAP rebinding can explain local extension after replication

To identify processes that could explain the 4D-MaxEnt-inferred changes in chromosome extension, we focus on two biophysical mechanisms that are known to affect bacterial chromosome compaction: NAPs and DNA supercoiling (Fig. 4B). First, we consider whether NAP redistribution during replication could locally extend newly replicated genomic regions. NAPs are highly abundant proteins that have been shown to compact DNA by up to 50% *in vitro* [43], although their role in bacterial chromosome condensation *in vivo* remains unclear. NAP dynamics during replication can be modeled using the Replication-Eviction (RE) model [31], which supposes that NAPs detach during replication, so that newly replicated chromosome regions have a lower NAP occupancy. We now extend this with a Replication-Eviction-Decompaction (RED) model: before NAPs rebind and compact newly replicated chromosomal regions, these regions will be extended compared to the rest of the chromosome. If NAP-mediated compaction equilibrates with a single time-scale after replication, this RED model predicts an exponentially decaying extension of newly replicated chromosomal regions, as inferred using the 4D-MaxEnt model (Fig. 4C).

On what time-scales would we expect NAP unbinding to locally extend newly replicated genomic regions? Since NAP diffusion is significantly faster than replication (HU diffuses across the cell in 10-40 seconds, based on a diffusivity of 0.1 −0.4 µm^2^s^*−*1^ [44]), evicted NAPs can rebind anywhere along the chromosome. This suggests that the NAP occupancy on newly replicated chromosomal regions should return to pre-replication levels at the rate of NAP rebinding. *In vivo* experiments suggest that these time-scales range from ∼0.1 s for HU [44] to ∼110 min for GapR [31]. Once NAPs have rebound on newly replicated DNA, the protein-DNA complex may still need to undergo conformational changes before final levels of compaction are reached. *In vitro*, DNA compacts on time-scales up to of order 60 min after an increase in HU concentration [45, 46]. This time-scale may increase with the degree of compaction, which is higher *in vitro* (∼50%) than we infer (∼10%). All in all, this suggests that NAP-mediated compaction might be re-established within an hour after a genomic region’s replication, consistent with the 10-20 minute time-scale inferred using the 4D-MaxEnt model. We hence conclude that a RED model could explain the local extension of newly replicated chromosomal regions inferred using the 4D-MaxEnt model.

### Positive supercoiling could cause compaction ahead of replication forks

Our 4D-MaxEnt model infers an approximately constant level of local compaction ahead of the replication forks. In these regions, we do not expect a change in NAP-mediated compaction; instead, the inferred compaction could be caused by increased levels of supercoiling.

Studies have demonstrated that both over- and under-winding (i.e. positive and negative supercoiling) of DNA lead to plectoneme formation and DNA compaction *in vitro* [47–49] and *in vivo* [50]. Whenever a replisome does not rotate as it progresses along the DNA helix, it pushes positive supercoils ahead of itself. This excess supercoiling is removed by topoisomerases, most notably gyrases, to prevent replication stalling [32, 33]. Both due to the rate of supercoil diffusion [51](Materials and Methods) and the presence of topological barriers approximately every 10 kb [14], we expect positive supercoiling to accumulate in a small region ahead of the replication forks. Once the activity of gyrases recruited to the replication forks is sufficient to counter-act positive supercoiling due to replication, these excess positive supercoiling levels should equilibrate, consistent with the approximately constant level of fork-associated compaction we infer for *t* ≥ 10 min (Fig. 4D). Hence, localized excess supercoiling could explain the compaction ahead of the replication forks inferred using the 4D-MaxEnt approach.

The inferred magnitude of relative extension ahead of the replication forks is also consistent with a supercoiling-based mechanism. Inhibition of gyrases, which maintain the chromosome in a negatively supercoiled state, has been shown to increase nucleoid volume by around 5% [52]. Simulation models, on the other hand, suggest that native levels of supercoiling in bacteria lead to a reduction of the radius of gyration of a DNA segment by around 8% [53]. These estimates are the same order of magnitude as the inferred ∼5% local compression ahead of the replication forks. Supercoiling is hence a plausible mechanism for establishing our inferred pattern of local compaction ahead of the replication forks.

### Incorporating replication-induced local extension changes into the mechanistic model

Finally, we construct a bottom-up model that captures both the large-scale organization and the local extension changes of the chromosome inferred using the 4D-MaxEnt method. Since supercoiling and NAP binding compact DNA on length-scales below 2.5 kb, the length corresponding to one monomer in our mechanistic model, our coarse-grained simulations are not suitable for explicitly modeling these mechanisms. However, a local change in DNA properties below the monomer length-scale can be effectively captured by locally modifying both the rest length and the stiffness of the corresponding spring (Materials and Methods). By including a replication-induced transient modification of the spring parameters both behind and ahead of the replication forks in our mechanistic model, we can capture the pattern of local extension and compression inferred using the 4D-MaxEnt approach (Materials and Methods, Fig. 5A).

**FIG. 5.**
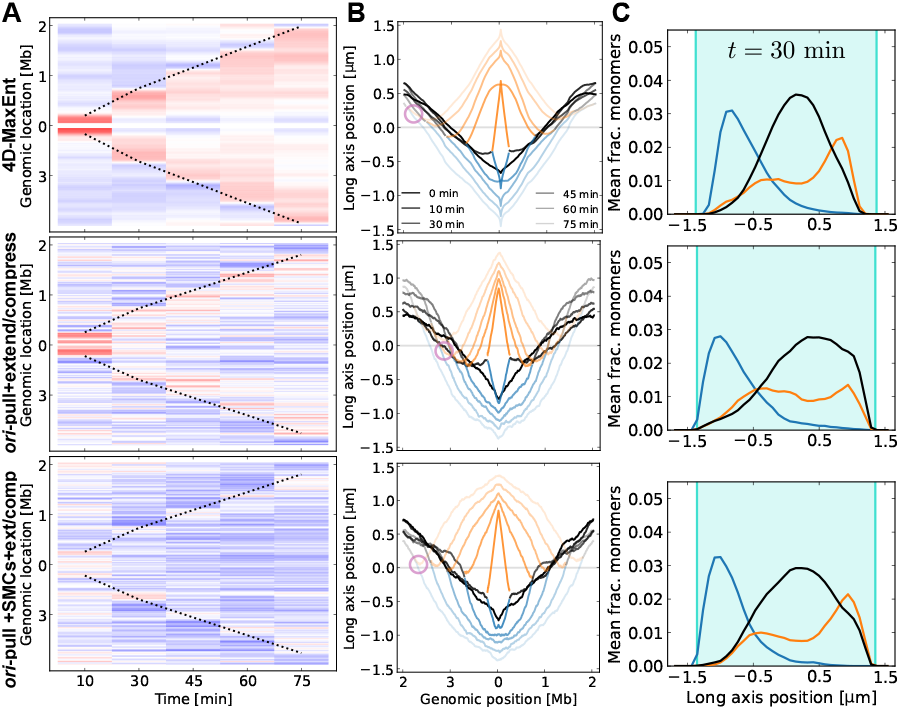
Mechanistic model with local extension and compression. Rows correspond to 4D-MaxEnt model; mechanistic model with *ori*-pulling together with extension behind and compression ahead of replication forks; mechanistic model with *ori*-pulling and 40 loop-extruders together with extension behind and compression ahead of replication forks. (A) Relative extension kymographs. Color bar in Fig. 4A. Mechanistic model with loop-extruders (bottom row) shows additional compaction due to the increased density of small loops during replication, as discussed in SI Appendix.(B) Mean long-axis positions of all loci. Circles show examples where newly replicated chromosomal regions have a smaller separation. (C) Densities of chromosomal strands across the confinement length at *t* = 30 min.

Despite significant changes in the local extensions of the polymer, *ori*-pulling still establishes linear organization of the chromosome in this full mechanistic model (Fig. 5B). Interestingly, after addition of local extension behind the replication forks, the mechanistic model predicts a smaller separation between newly replicated chromosomal regions, consistent with 4D-MaxEnt inferences (circles in Fig. 5B; compare to Fig. 3B,C; SI Appendix Fig. S13B). The closer association of the replicated strands is also reflected in the density histogram of the *ter*-proximal strand, which now shows a bimodal distribution, as seen in the 4D-MaxEnt model (Fig. 5C).

We also find that loop-extruders are still necessary to segregate the unreplicated chromosome regions to mid-cell by *t* = 30 min (Fig. 5C). Taken together, our full mechanistic model captures all the main inferences made using the 4D-MaxEnt method, and hence sheds light on the underlying mechanisms of bacterial chromosome organization.

## DISCUSSION

We have introduced a fully data-driven model for the organization of a replicating bacterial chromosome. Our approach distinguishes itself from previous works [21, 22] by introducing a principled, unbiased model selection method that integrates time-course Hi-C with fluorescence microscopy data and accounts for factors such as cell growth and the indistinguishability of replicated chromosomal strands in Hi-C reads. Using independent microscopy experiments, we confirm that our model accurately infers single-cell 3D chromosome conformations, and their inherent variability, across the *C. crescentus* cell cycle.

In addition to providing a minimal physical interpretation of time-course Hi-C data, our data-driven model can be used to develop a mechanistic understanding of chromosome organization. We used our 4D-MaxEnt model to constrain a minimal mechanistic model of a replicating bacterial chromosome. We find that, although a ParAB*S* -like *ori*-pulling force is sufficient for establishing linear chromosome order during replication, loop-extruders are required to ensure the timely localization of terminal regions to mid-cell. Since loop-extruders in our model are mainly found within 600 kb of the origins of replication, based on experiments [8], they affect the localization of terminal regions indirectly, by introducing constraints that alter the chromosome’s effective topology and compaction [11, 54]. Since *C. crescentus* Δ*smc* mutants do not show significant segregation defects [8, 14, 31], the segregation of terminal regions might also be orchestrated by other late-acting mechanisms, such as the attachment of the terminal regions to the divisome by ZapT [55], and active terminus-segregation by the translocase FtsK [56].

The local compaction of bacterial chromosomes is intrinsically connected with transcriptional processes [57, 58], but it remains unclear how replication affects this interplay. Our 4D-MaxEnt model reveals chromosome compaction level changes by up to 60% around the replication forks, lasting for tens of minutes, a significant amount of time compared to the *C. crescentus* cell cycle. We propose that these inferred local extension patterns would be consistent with NAP unbinding behind and positive supercoiling ahead of the replication forks. Although we considered these two mechanisms separately, NAP binding is known to be affected by supercoiling levels [59–61], and plectonemes can be stabilized by NAPs [62]. Therefore, synergistic effects between NAP binding and supercoiling could also contribute to chromosome compaction dynamics. We speculate that chromosome decompaction behind the replication forks might improve the local accessibility of the chromosome, and thereby also affect transcription levels [5, 58].

Time-course Hi-C data contain information about chromosome organization both across genomic scales and across the cell cycle in both pro- and eukaryotic systems [63]. Although such data are still limited in availability, methods to synchronize bacterial populations– such as treatment with serine hydroxamate [64], depletion of DnaA [65], or use of a “baby machine” [66]–do exist. Given time-course data, extracting information about the dynamics of the system in a faithful way is still challenging. Here, we used an information theory-based approach to construct a rigorous model inference scheme for time-course Hi-C data across the bacterial cell cycle. This approach may serve as a starting point for constructing similarly rigorous models for more complex systems, such as fast-growing bacteria undergoing multi-fork replication or eukaryotes.

## MATERIALS AND METHODS

### Processing of Hi-C data

Input Hi-C maps from [14] were first used to calculate the expected replication fork positions and were then normalized (SI Appendix, Fig. S1). For *t* = 0, each column of the Hi-C matrix was normalized to sum to 1 [14].

For later time-points, the Hi-C count for each bin was divided by the count at *t* = 0. This gives a corrected normalization factor taking into account that replicated regions form more contacts than unreplicated ones.

### Imaging data

To follow chromosome dynamics in live cells, we imaged synchronized *C. crescentus* cells, which expressed the fluorescent proteins, TetR-YFP and LacI-CFP, binding to tetO and lacO arrays introduced at specific chromosomal loci. For this, we used previously constructed strains [30, 38](SI Appendix, Table S1). Following the procedure described in [14], we synchronized *C. crescentus* swarmer cells (see SI Appendix for more details), which initially contain a single, unreplicated chromosome. Cells were then imaged via fluorescence microscopy (see SI Appendix for further details) at all cell cycle time-points for which Hi-C data was available [14]. The MicrobeJ plugin [67] for ImageJ [68] was used to segment the cells, after which parameters such as inter-focal distances and cell dimensions could be measured.

For each time-point, we measured the mean length of the cells and used this value minus the estimated cell envelope thickness of 61 nm [69] as the 4D-MaxEnt model’s confinement length. To obtain the constraints on the mean *ori* distance for each replication stage, we measured the mean distance between two fluorescently labelled *ori* loci at each Hi-C time-point (Fig.2C, (SI Appendix). The experimental data used for model validation (Fig.2B) were obtained similarly, using strains with different foci labeled (SI Appendix, Table S1).

### Chromosome density histograms

To visualize patterns in chromosome segregation, we calculate histograms that show the relative density of monomers belonging to different DNA strands across the long axis of the cell. Briefly, in 4D-MaxEnt lattice polymer simulations, we calculate the number of monomers at a given discrete long-axis position for each strand. This way, we find three unnormalized histograms, corresponding to the two replicated strands and the unreplicated strand of the chromosome. To normalize the histograms, we divide all counts by the total number of replicated monomers.

For mechanistic MD simulations conducted off-lattice, we bin the long-axis positions of the monomers, using bins that correspond to the 4D-MaxEnt model’s discretized long axis positions. We note that since the mechanistic simulations are conducted off-lattice, the accessible volume near the confinement volume boundaries is higher than in the 4D-MaxEnt model, which could explain some of the differences between histograms and mean long-axis positions of the models (Fig. 3D-F).

### Mechanistic model with *ori*-pulling and loop extrusion

Molecular dynamics simulations to model chromosome segregation were conducted as in [11, 12], based on earlier simulations using the polychrom wrapper for OpenMM [41, 42] and the looplib package for loop extrusion simulations [70] (see SI Appendix for details).

In loop-extruder simulations, we assume that *M*_0_ bidirectional loop-extruders per chromosome, with a processivity of 1200 kb and a leg speed of 18 kb per minute [8], are loaded primarily at the origin of replication [71]. We vary the total number of monomers in our simulations, *N*, and find similar large-scale dynamics of the chromosome (SI Appendix, Fig. S14, S15). We also run our simulations for different values of *M*_0_, and find that using approximately *M*_0_ = 40 loop-extruders per chromosome gives organization similar to the 4D-MaxEnt model (SI Appendix, Fig. S9), consistent with previously used values for *B. subtilis* [71]. Unless otherwise stated, results are shown for *M*_0_ = 40 and *N* = 1620, the same coarse-graining scale as in the 4D-MaxEnt model.

Briefly, we first simulate loop extrusion on a 1D lattice corresponding to the replicating chromosome. Two replication forks move forward independently. Upon collisions between replication forks and loop-extruders, we assume that loop-extruders unbind. To keep the loop-extruder density on the chromosome constant, the number of loop-extruders is increased during replication. The initial positions of the first *M*_0_ loop-extruders are taken from the final state of the previous simulation, to model the effects of continuous growth in loop-extruder numbers (SI Appendix, Fig. S16). Loop-extruders bind at the *ori* region with a 50% probability, and have a factor 100 times higher off-loading rate near the terminus. Upon head-on collisions between loop-extruders, the motor proteins stall, unbind, or bypass each other [71].

For each set of parameters, we save 200 1D loop extrusion trajectories. The loop-extruder and replication fork positions are then used to constrain molecular dynamics simulations. We simulate a replicating bead-spring polymer in a growing cylindrical confinement with rounded caps, with loop-extruders modeled as additional, moving springs between monomers. *ori*-pulling is modeled by constraining the long-axis positions of each origin of replication using a harmonic potential. The distance between the potentials’ minima is set by a function fitted to the measured *ori-ori* distances used to constrain the 4D-MaxEnt model [11].

For each coarse-graining scale, we adjust the spring length *ℓ*_0_ and the spring coefficient *k* so that the mean and standard deviation of extensions of 10 kb regions match the 4D-MaxEnt model (SI Appendix, Fig. S14 A-D). The time-scale of the simulations is set to be consistent with mean-squared displacement data for *E. coli* loci at short time-scales [72] (SI Appendix, Fig. S14E).

### Local extension is inferred from Hi-C constraints

The 4D-MaxEnt model infers persistent local extension behind and compaction ahead of the replication forks. Since the *ori*-constrained model without Hi-C constraints shows close to zero relative extension throughout the replication cycle (SI Appendix, Fig. S11A), Hi-C constraints seem necessary to infer local changes in chromosome compaction. We also analyze local extensions of the model with only Hi-C constraints, but no *ori-ori* distance constraints. This unsegregating model also shows local extension behind and compaction ahead of the replication forks (SI Appendix, Fig. S11A), although extension behind the forks does decays slower than in the 4D-MaxEnt model. We hence conclude that the inferred local extension pattern arises from the imposed Hi-C constraints.

### Estimating supercoil diffusion ahead of replication forks

In single molecule studies of plectoneme diffusion on a 21 kb DNA segment held under tension, it was found that plectonemes diffuse either continuously, or by large jumps [51]. The mean diffusion constant for continuous trajectories was found to be of order *D*_c_ ≈0.01 −0.1 µm^2^s^*−*1^, depending on the tension exerted on the DNA. On the other hand, hops of size 0.1−1 µm were observed, with a mean hopping time of order 0.1 s. Assuming a random walk of hops, the hopping diffusivity would be *D*_h_ ≈ 0.1 − 10 µm^2^s^*−*1^.

Assuming a hopping distance of 0.1 μm, and an ex-tension of of 5 µm for 15 kb in the experiments [51], we would find *D*_h_ ≈0.9 kb^2^s^*−*1^, meaning that plectonemes would diffuse through a CID of length 100 kb in hours,whereas the region would be replicated in roughly 4 minutes. Since CID boundaries correspond to highly transcribed genes that act act as supercoiling diffusion barriers [57], we would expect supercoiling to accumulate ahead of the replication forks, as discussed in the main text.

Assuming longer hops of size 1 µm, we would find *D*_h_ ≈90 kb^2^s^*−*1^, meaning that plectonemes would diffuse through a CID of length 100 kb in two minutes, similar to the time-scale of the CID’s replication. In this case, supercoiling would still accumulate in a region ahead of the replication forks, but they would be distributed over a broader region due to diffusion. If supercoil diffusion is restricted to 10 kb domains [14], we would also expect positive supercoiling to accumulate in a small region ahead of the replication forks.

### Modeling changes in local extension around the replication forks

Any changes in the contour length *ℓ*_c_ and/or persistence length *ℓ*_p_ of a segment of the chromosome represented by one bead/spring in our mechanistic model will translate to a change in both the coarse-grained spring’s rest length and stiffness. The radius of gyration of the DNA segment (the spring rest length) scales as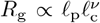 [73], where *ν* is the Flory exponent. The stiffness of this entropic spring, on the other hand, scales as 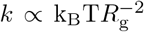. Hence, to model changes in local extension in our mechanistic model, we locally adjust the spring stiffnesses, spring rest-lengths and excluded volume radii ahead and/or behind the replication forks. If the rest length and excluded volume radius of a spring is adjusted by a factor *x*, the corresponding spring stiffness is adjusted by a factor *x*^*−*2^.

Motivated by the 4D-MaxEnt model’s local extension patterns, for extension behind the forks we modify the spring rest length *ℓ* assuming an exponential decay with the distance to the closest fork ahead of a replicated monomer, *s*:

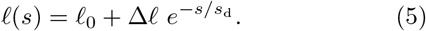

Here *ℓ*_0_ is the original spring rest length, Δ*ℓ* = 0.1*ℓ*_0_ for extension, and *s*_d_ = 600 kb is the decay length. The value for Δ*ℓ* was motivated by the initial extension of 10% seen in the 4D-MaxEnt model, and the length-scale *s*_*d*_ was similarly motivated by the observed extensions over length scales of up to 1 Mb (Fig. 4A). For simulations with compression ahead of the replication forks, *s* is the distance to the nearest fork *behind* the given monomer, Δ*ℓ* = −0.1*ℓ*_0_, and *s*_d_ = 100 kb. We assume that local compression begins 5 minutes after the start of replication, to ensure that at *t* = 0 min the chromosome is uncompressed. We find that using only extension behind the forks does not cause compression ahead of the forks, and vice versa (SI Appendix, Fig. S13). The addition of temporary compaction and extension slightly changes the frequencies of contacts between fork-proximal genomic regions (SI Appendix, Fig. S17).

## ACKNOWLEDGMENTS

We thank the labs of Lucy Shapiro and Patrick Viollier for sharing *C. crescentus* strains. We also thank Suckjoon Jun, Remus Dame, Capucine Beraud, and members of our group for fruitful discussions and feedback. This project has received funding from the European Research Council (ERC) under the European Union’s Horizon 2020 research and innovation programme (Grant agreement No. 101122863), and from the Max Planck Society under the Max Planck Fellowship.

## CODE AND DATA AVAILABILITY

Raw simulated data is at 10.5281/zenodo.15099819. 4D-MaxEnt simulation code is at github.com/JorisJBM/ChromMaxEnt4D. Mechanistic simulation code is at github.com/PLSysGitHub/loop-extrusion_with_replication. Analysis code is at github.com/PLSysGitHub/4D_MaxEnt_analysis.

