## Supplementary information for "Learning the dynamic organization of a replicating bacterial chromosome from time-course Hi-C data"

**3 Deriving the form of the 4D-MaxEnt model** **5**

**4 Monte Carlo simulations** **6**

**5 Model validation** **7**
- 5.1 Correcting for indistinguishability of fluorescent foci at short distance . 7

**8 Supplementary references** **33**

### 1 Experimental procedures on *C. crescentus* cells

#### 1.1 Bacterial strains and growth conditions

The *C. crescentus* strains used in this study (Table S1) were derived from the synchronizable wild-type CB15N (NA1000). Cells were grown in peptone-yeast extract (PYE) medium (Pointdexter, 1964) at 28°C under aerobic conditions (shaking at 190 rpm).

#### 1.2 Synchronization of *C. crescentus* cultures

In order to analyze *C. crescentus* cells in a specific phase of their cell cycle, corresponding to a specific stage in their replication and segregation process, we synchronized the cells according to the protocol established in [1]. In brief, *C. crescentus* cells were grown to early exponential phase (OD $\sim$0.1) in PYE and induced for 2h with 2 µM xylose in order to express YFP and CFP that bind their respective arrays at specific chromosomal loci (see Table S1). Afterwards, cells were pelleted, resuspended in M2 salts buffer [2] and mixed 1:1 with Percoll. In a density centrifugation step, the newborn swarmer cells were separated from the stalked cells. The swarmer cells were collected and washed once in M2 salts, before being released into PYE (including 2 µM xylose) and allowed to grow at 28°C until they were analyzed by microscopy.

#### 1.3 Experimental determination of cell sizes and intracellular locations of chromosomal loci throughout the cell cycle

To determine the dimensions of *C. crescentus* cells, as well as the copy number and intracellular location of specific fluorescently labeled chromosomal loci at specific time points in their cell cycle, we subjected cells at specific time points after synchronization (see above) to fluorescence microscopy. To this end, cells were immobilized on pads made of 1% agarose in water and observed with a Nikon Ti2 Eclipse microscope. The microscope was equipped with an alpha Plan Apo $\lambda$ 100x/1.45 Oil ($\infty$)/0.17 WD 0.13 Ph3 objective (Nikon, Japan), a Spectra X Light Engine (Lumencor, USA) light source and a CFP-2432C and a YFP-2427B filter (Semrock, USA). Images were collected with an Orca-flash4.0LT Plus C11440-42U30 camera (Hamamatsu, Japan) and recorded with NIS Elements 5.30.02 (Nikon, Japan).

In order to extract the cellular dimensions and intracellular positions of the fluorescently-labeled loci, cells were segmented based on the phase contrast channel using the ImageJ [3] plugin MicrobeJ [4]. To be able to monitor cell cycle progression in each subset, both cell lengths and the percentage of cells showing constrictions (as detected via MicrobeJ's feature constriction option) were followed for each time-point after synchronization for each strain. Fluorescent maxima were detected using the maxima detection, which determines the localizations of the maxima relative to the poles of the cell at sub-pixel resolution using a Gaussian fit.

#### 2 Model input data

For each time-point that we model, we need the following data:

1. mean distances between replicated *ori*s
2. bias-corrected Hi-C data,
3. the mean replication fork position,
4. the mean cell length,

The mean separations between *ori*s were obtained as described in the last section. We now discuss how the rest of the input data was obtained and processed.

##### 2.1 Input Hi-C data

As input data, we used the synchronized Hi-C data for *C. crescentus* cells, collected 0, 10, 30, 45, 60, and 75 minutes after replication initiation [5]. These time-points define the temporal resolution of our model.

The raw Hi-C counts $m_{ij}(t)$ cannot be used directly, since they are biased due to varying crosslinker affinities between genomic regions, and to a lesser extent differences in GC content and the mappability of individual reads [5]. To remove such biases for the data set at $t = 0$, we used the same procedure as in [5]: the data is normalized so that each row and column of the Hi-C map sums to one; the fraction of corrected counts $m_{ij}^{\text{cor}}$ in each column should hence be $f_i^{\text{corr}}(0) = \frac{\sum_j m_{ij}^{\text{cor}}}{\sum_{k,l} m_{kl}^{\text{cor}}} = \frac{1}{N_{\text{bins}}}$, where $N_{\text{bins}}$ is the number of Hi-C bins. This normalization procedure assumes that all genomic regions have an equal number of expected contacts.

To normalize the Hi-C maps for later time-points, we make use of the column sums in the raw data set at $t = 0$. The fraction of raw contact counts in column $i$ at $t = 0$, $f_i(t = 0)$, reflects the read count bias of site $i$. Thus, for each data set at a later time $t > 0$, we define the bias-corrected column fraction as $f_i^{\text{corr}}(t) = \frac{f_i(t)}{f_i(0) \sum_j f_j(t)/f_j(0)}$ (Supplementary Figure S2). Setting $f_i^{\text{corr}}$ as the target column fraction assumes that biases in crosslinker affinities and the mappability of individual reads remain constant throughout the cell cycle.

For a given time-point $t$, to construct an input contact frequency data set $f_{ij}^{\text{expt}}(t)$ where the column fractions match $f_i^{\text{corr}}$, we use an iterative procedure [5]. For the initial iteration step, we set $\tilde{m}_{ij}(t) = m_{ij}(t)$ and then calculate the current column fractions

$\tilde{f}_i(t) = \sum_j \tilde{m}_{ij}(t)/\tilde{S}(t)$, where $\tilde{S}(t) = \sum_{i,j} \tilde{m}_{ij}$. Each Hi-C matrix entry $\tilde{m}_{ij}(t)$ is then rescaled according to $\tilde{m}_{ij}(t) \to \tilde{m}_{ij}(t)\frac{f_i^{\mathrm{corr}} f_j^{\mathrm{corr}}}{\tilde{f}_i(t)\tilde{f}_j(t)}$. We then recalculate $\tilde{f}_i$, and repeat the process until the target column fractions are matched within an average relative deviation of 1 in 100000. Lastly, the entire Hi-C matrix is rescaled such that the sum of all entries equals the number of columns. The resulting input Hi-C maps $f_{ij}^{\mathrm{expt}}(t)$ for each cell cycle time $t$ are shown in Supplementary Figure S1.

##### 2.2 Replication fork positions

To estimate the mean replication fork position for each Hi-C data set following [5], we use the bias-corrected column sums $\sigma_i^{\mathrm{corr}}$. Briefly, replicated loci should be roughly two times likelier to be present in Hi-C reads than unreplicated ones. For each replication stage, we hence calculate the relative number of Hi-C reads compared to $t = 0$ for all loci (Supplementary Figure S2A). The inflection points of the transition from high relative counts around the *ori* towards low relative counts around the *ter* are taken as the mean replication fork positions (Supplementary Figure S2B). These inflection points are found by smoothing the relative contact counts and taking a numerical derivative.

Combining the estimated mean replication fork positions for the Hi-C data sets at 30, 45, and 60 minutes after synchronization, we find that the inferred locations lie closely along a linear fit for both chromosomal arms (Supplementary Figure S2C). Importantly, the replication progress for both chromosomal arms is determined independently. Thus the close agreement between the replication speeds on the two arms confirms the accuracy of the inferred replication progress. The mean speeds of the linear fits for both arms are used to define the fork positions at each time-point in the 4D-MaxEnt model.

Replication fork speed has been suggested to depend on transcriptional activity, and to slow down near highly expressed genes [6]. This could explain why we infer slower fork progression between 60 and 75 minutes.

##### 2.3 Cell size

To determine the mean cell size associated with each replication fork position, we make use of the fluorescent microscopy images gathered for calculating locus distances. For each time-point, the mean cell length is determined (blue dots in Supplementary Figure S3A). From this, the estimated cell envelope width of 61 nm [7, 8] is subtracted, yielding the confinement lengths used as model inputs (black dots in Supplementary Figure S3A). For the confinement width, a cylinder with rounded caps as used in [8] is applied, with the confinement width of 0.63 µm assumed to be constant throughout the cell cycle [9, 10].

#### 3 Deriving the form of the 4D-MaxEnt model

To derive the functional form of the 4D-MaxEnt model, we must maximize the Shannon Entropy after applying the appropriate constraints. We define $f_{i,j}$ as the combined number of *cis*- and *trans*-contacts between genomic locations $i$ and $j$ for a given lattice polymer configuration $\{\mathbf{r}, \mathbf{r}'\}$:

$$f_{i,j}(\{\mathbf{r}, \mathbf{r}'\}, t) = \begin{cases} \delta_{\mathbf{r}_i,\mathbf{r}_j}, \text{if } i \text{ and } j \text{ unreplicated,} \\ \delta_{\mathbf{r}_i,\mathbf{r}_j} + \delta_{\mathbf{r}_i,\mathbf{r}'_j}, \text{if } i \text{ unreplicated,} \\ \delta_{\mathbf{r}_i,\mathbf{r}_j} + \delta_{\mathbf{r}'_i,\mathbf{r}_j}, \text{if } j \text{ unreplicated,} \\ \delta_{\mathbf{r}_i,\mathbf{r}_j} + \delta_{\mathbf{r}'_i,\mathbf{r}'_j} + \delta_{\mathbf{r}_i,\mathbf{r}'_j} + \delta_{\mathbf{r}'_i,\mathbf{r}_j}, \text{if both replicated.} \end{cases} \tag{S1}$$

Our condition is that the mean of $f_{i,j}$ at a given time-point $t$ should be proportional to the Hi-C score $H_{i,j}(t)$:

$$\langle f_{i,j}(t)\rangle = \sum_{\{\mathbf{r},\mathbf{r}'\}} P(\{\mathbf{r}, \mathbf{r}'\}, t) f_{i,j}(\{\mathbf{r}, \mathbf{r}'\}, t) \stackrel{!}{=} C(t) H_{i,j}(t), \tag{S2}$$

Here $C(t)$ is the unknown scaling coefficient between the contact frequencies and the Hi-C counts. This scaling coefficient can differ between Hi-C maps for different replication stages.

In addition to the Hi-C constraints, we apply a constraint on the mean long-axis separation between origins of replication, $d_{\text{ori}}(\{\mathbf{r}, \mathbf{r}'\}) = |r^z_1 - r'^z_1|$. We thus impose:

$$\langle d_{\text{ori}}(t)\rangle = \sum_{\{\mathbf{r},\mathbf{r}'\}} P(\{\mathbf{r}, \mathbf{r}'\}, t) d_{\text{ori}}(\{\mathbf{r}, \mathbf{r}'\}) \stackrel{!}{=} \langle d^{\text{expt}}_{\text{ori}}(t)\rangle, \tag{S3}$$

where $\langle d^{\text{expt}}_{\text{ori}}(t)\rangle$ is the experimentally measured mean long-axis *ori* distance.

Finally, we require that the distribution is normalized:

$$\sum_{\{\mathbf{r},\mathbf{r}'\}} P(\{\mathbf{r}, \mathbf{r}'\}, t) \stackrel{!}{=} 1. \tag{S4}$$

To maximize the distribution entropy under constraints (S2), (S3), and (S4), we introduce the functional $\tilde{S}[P(\{\mathbf{r}, \mathbf{r}'\}, t)]$, with Lagrange multipliers $\epsilon_{i,j}(t)$ for each Hi-C constraint, $\alpha(t)$ enforcing the separation between *ori*'s, and $\lambda(t)$ ensuring normalization:

$$
\begin{aligned}
\tilde{S}[P(\{\mathbf{r},\mathbf{r}'\},t)] = &- \sum_{\{\mathbf{r},\mathbf{r}'\}} P(\{\mathbf{r},\mathbf{r}'\},t) \ln P(\{\mathbf{r},\mathbf{r}'\},t) \\
&- \sum_{i\leq j} \epsilon_{i,j}(t) \Bigg( \sum_{\{\mathbf{r},\mathbf{r}'\}} P(\{\mathbf{r},\mathbf{r}'\},t) f_{i,j}(\{\mathbf{r},\mathbf{r}'\},t) - C(t) M_{i,j}(t) \Bigg) \\
&- \alpha(t) \Bigg( \sum_{\{\mathbf{r},\mathbf{r}'\}} P(\{\mathbf{r},\mathbf{r}'\},t) d_{\mathrm{ori}}(\{\mathbf{r},\mathbf{r}'\}) - \langle d^{\mathrm{expt}}_{\mathrm{ori}}(t) \rangle \Bigg) \\
&- \lambda(t) \Bigg( \sum_{\{\mathbf{r},\mathbf{r}'\}} P(\{\mathbf{r},\mathbf{r}'\},t) - 1 \Bigg).
\end{aligned} \tag{S5}
$$

Here, all sums over $i, j$ run for unique pairs of monomers with $i \leq j$. Setting $\frac{\partial \tilde{S}}{\partial P(\{\mathbf{r},\mathbf{r}'\},t)} \stackrel{!}{=} 0$ we obtain:

$$
P(\{\mathbf{r},\mathbf{r}'\},t) = \frac{1}{Z} \exp\Big[ -\alpha(t) d_{\mathrm{ori}}(\{\mathbf{r},\mathbf{r}'\}) - \sum_{i\leq j} \epsilon_{i,j}(t) f_{i,j}(\{\mathbf{r},\mathbf{r}'\},t) \Big], \tag{S6}
$$

where $Z = \exp[1+\lambda]$. Finally, maximizing (S5) with respect to the scaling coefficient $C(t)$ gives rise to another constraint:

$$
\sum_{i,j} \epsilon_{i,j}(t) M_{i,j}(t) = 0. \tag{S7}
$$

For a converged model, which satisfies (S2), this is equivalent to a condition on the total contact energy of the system:

$$
\Bigg\langle \sum_{i,j} \epsilon_{i,j}(t) f_{i,j}(t) \Bigg\rangle = 0. \tag{S8}
$$

(S6) defines the 4D-MaxEnt model. The effective energies $\epsilon$ and $\alpha$ must be found such that (S2), (S3) and (S8) are satisfied. Analogously to [8], a solution to this large set of simultaneous equations can be found by mapping Eq. S6 to an equilibrium polymer model. This equilibrium polymer model contains close-range pairwise interaction energies $\epsilon_{ij}(t)$ between monomers, and a pushing force $\alpha(t)$ that couples to the distance between replicated *ori*'s.

### 4 Monte Carlo simulations

The Monte Carlo simulation of the lattice polymer is performed as in [8], with one extra move included to change the replication fork position. The added 'fork move', illustrated in Supplementary Figure S3B, together with the loop move, kink move, and crankshaft move [11], forms the set of moves randomly chosen from at each step of the Monte Carlo simulation.

The addition of the fork moves preserves ergodicity, which we can see as follows. The unreplicated strand and one of the replicated strands, (for example the *ter*-distal strand), together have the topology of a single unreplicated chromosome. Except for the replication fork sites $F_1, F_2$, this subset $S$ of monomers can be modified in the same way as the unreplicated chromosome in [8], which was shown to be ergodic under the loop, kink, and crankshaft moves. The replication fork sites $F_i$ are only modified by the fork move. From the perspective of subset $S$ however, the fork move simply behaves as either a kink move or a loop move, with the constraint that the first monomer of Strand 2 is in the right position to allow these moves.

This constraint is satisfied as long as it is possible to put the first monomer of Strand 2 at each of the six possible sites around the replication fork site by a sequence of polymer moves. Here, Strand 2 can be considered as a linear polymer with fixed endpoints, where each of its monomers is subject to the kink, loop and crankshaft moves. Thus, Strand 2 is subject to ergodic sampling. The only case where it is not possible to put the first monomer of Strand 2 at any position around the replication fork is if the replicated strand is completely stretched out, i.e. the distance between the replication forks is equal to $F_1 + N - F_2$. Since for all sampled time-points, the contour length of the replicated region is longer than our confinement length, this exception is not relevant for our model.

As the crankshaft move can be constructed as a combination of loop and kink moves, the replication fork site $F_i$ is effectively subject to the same moves as the other monomers in subset $S$, and thus the monomers in subset $S$ are subject to an ergodic sampling of states. As the *ter*-distal and the *ter*-proximal strand are identical, both possible subsets that can be constructed are subject to ergodic sampling, thus this also holds for the chromosome as a whole.

### 5 Model validation

#### 5.1 Correcting for indistinguishability of fluorescent foci at short distance

For comparison of 4D-MaxEnt inferences to validation data, we perform a conditional averaging on simulated localizations to directly compare experimental data with model inferences. This conditional averaging is required because after a chromosomal region has been replicated, the 4D-MaxEnt model always contains exactly two copies of this region, whereas either one or two copies are typically observed in experiments. This discrepancy is attributed to a combination of imperfect synchronization of cells and imperfect label detection. One contribution to the imperfect label detection is the indistinguishability of fluorescent foci at short distance, which results in a systematic bias in inferred mean label positions. In this section we estimate the magnitude of this bias, and describe how we compute the bias-corrected localizations for the 4D-MaxEnt model.

To estimate the distance at which fluorescent foci become indistinguishable, we construct a histogram of the experimentally observed pairwise focus distances (Supplementary Figure S5A). In this histogram, we find a sudden cutoff to zero counts for distances below 0.32 $\mu$m. This cutoff is consistent with our assumption of indistinguishability of fluorescent foci at short distance. The observed minimum distance of 0.32 $\mu$m is used as a cutoff value in the calculation of bias-corrected 4D-MaxEnt localizations.

In Supplementary Figure S5B, the measured fractions of cells with two foci are shown for all measurement conditions, together with the value inferred using the 4D-MaxEnt method, given the indistinguishability of foci at short distance. The results suggest that this indistinguishability strongly contributes to the observed fractions. Furthermore, we find that the onset of replication for tagged regions, taken as the onset of non-zero fractions, matches well between model and experiment.

To calculate the bias-corrected 4D-MaxEnt localization profiles shown in Main Text Fig. 2, we generate an ensemble of 4D-MaxEnt chromosome configurations for each time point. For each configuration, we compute 2D distances between the experimentally tagged regions, projected along the cell length and cell width coordinates. Only loci pairs whose projected 2D distance is above the resolution limit are included to obtain the bias-corrected localization profiles shown in Main Text Fig. 2. The same profiles without the bias correction are shown in Supplementary Figure S6. Without the correction, replicated focus pairs are inferred to be closer to each other shortly after replication.

##### 5.2 Ori-constrained model performs worse for validation data

As explained in the main text, we construct a maximum entropy model where only the distance between the *ori*s is constrained. This *ori*-constrained model does not infer the same linear organization or the density histograms as the Hi-C constrained 4D-MaxEnt model (Supplementary Figure S7A,B; compare to Main Fig. 3A and Supplementary Figure S8). Furthermore, it infers the locus positions used to validate the model less accurately than the 4D-MaxEnt model (Supplementary Figure S7C).

#### 6 Mechanistic model

##### 6.1 Model description

For 1D simulations of loop extrusion on a replicating bacterial chromosome, we add replication to our previous model [12] built on the looplib package [13]. To model replication, we include two replication forks moving independently in opposite directions. Replication forks create new lattice sites accessible to loop-extruders. For simplicity, when a replication fork encounters a loop-extruder, the loop-extruder unbinds. In previous work, we found that allowing loop-extruders to by-pass replication forks did not qualitatively change segregation dynamics [12].

The loop-extruder off-loading rate is set by the extrusion time $\tau = \lambda/(2v_{\mathrm{LE}})$ of the loop-extruders, determined by the processivity $\lambda$. We use values of $\lambda = 1200$ kb and $v_{\mathrm{LE}} = 18$ kb/min, in agreement with experimental data [14]. In a 10 kb region around the terminus, the off-loading rate is increased by a factor of 100. Loop-extruders have a 50% chance of being loaded at the origin of replication, as used in previous simulations [15]. We vary the number of loop-extruders per chromosome, $M_0$, to qualitatively match the 4D-MaxEnt model (Supplementary Figure S9), and find that $M_0 = 40$ gives a good fit, in agreement with previous simulation works that have used 40 loop-extruders per chromosome to model the *B. subtilis* chromosome [15]. Using fewer loop-extruders matches the mean-long axis curves of the 4D-MaxEnt better, but segregation of the unreplicated strand to mid-cell appears worse (Supplementary Figure S9C).

To keep the density of loop-extruders constant, the number of loop-extruders is increased during replication. In real cells, we expect the loop-extruder number to rise continuously, rather than starting suddenly at replication initiation. To model this, each 1D loop-extruder simulation "inherits" its initial loop-extruder positions from the final loop-extruder positions of the last simulation. After roughly 10 generations, the initial loop-extruder distributions have converged (Supplementary Figure S16). To ensure that the initial loop-extruder positions are sampled from the converged distribution, we hence skip the first 20 generations of 1D simulations.

For 3D polymer simulations, we use the polychrom wrapper [16] for the molecular dynamics library OpenMM [17]. A bead-spring polymer is confined to a cylinder with rounded caps using a harmonic potential. The cylinder's length grows in time at an exponential rate [12].

Unreplicated monomers have no excluded volume interactions, and are not connected by springs to any other monomers. When a replication fork moves from site $i$ to $i+1$, we initialize the position (velocity) of bead $(i+1)'$ (the replicate of $i+1$) as the mean of the positions (velocities) of beads $i+1$ and $i'$. We also turn on excluded volume interactions for monomer $(i+1)'$, and turn on the harmonic potential between neighboring monomers $[i', (i+1)']$. Additionally, we add a temporary harmonic potential between replicates $[i+1, (i+1)']$ at the new fork site. This additional force keeps the replicated strands tied together. It is turned off when the fork moves again.

Excluded volume interactions between replicated beads are modeled by a repulsive potential

$$U(r) = U_0 \left(1 + \left(\frac{r}{R_{\mathrm{max}}}\right)^{12} \left(\frac{6}{7}\left(\frac{r}{R_{\mathrm{max}}}\right)^2 - 1\right)\right), \tag{S9}$$

where $U_0 = 5\ \mathrm{k_BT}$ is the depth of the potential; $r$ is the separation between two beads; and $R_{\mathrm{max}}$ is the distance at which the potential's derivative becomes zero (1.05 times the spring length $b = 0.03$ µm). This potential allows occasional strand passing, mimicking the effect of topoisomerases *in vivo*, and our simulations hence do not conserve the chain topology. However, the potential is more than sufficient for linear chains to swell and show the appropriate Flory exponent of $\nu = 3/5$ [12].

A 3D simulation time-step consist of updating the replication fork positions, turning on interactions for newly replicated monomers, updating the positions of additional springs representing loop-extruders based on the 1D simulations, and then allowing the polymer to relax. For each set of simulation parameters, at least 200 simulations running over the entire replication cycle are conducted. All simulations are initialized from an unreplicated *ori-ter* configuration.

##### 6.2 Setting model length- and time-scales

In previous work, we modeled the segregation of a *C. crescentus*-like chromosome using a coarse-graining level where one spring in the simulations corresponded to 10 kb [12]. For this study, we mainly use beads corresponding to 2.5 kb, the same coarse-graining scale as the 4D-MaxEnt model. We also perform mechanistic *ori*-pulling simulations using $N = 404$, 1620, and 4050, to check that our results are not strongly dependent on the coarse-graining scale.

When adjusting the coarse-graining scale, we want to ensure that the physical properties and time-scales of the system do not change. Therefore, when changing the number of monomers from $N$ to $N'$, we make the following edits:

1. We choose the monomer size $b'$ and spring constant such that the mechanistic model matches the 4D-MaxEnt model's distribution of end-to-end distances for 10 kb region extensions (Supplementary Figure S14).
2. To keep the total mass of the chain constant, we change $m \to m\, N'/N$.
3. To keep the total friction of the chain in a Rouse-like setting constant, we change the friction coefficients by $\zeta \to \zeta\, N'/N$.

In the case of an ideal polymer, the above changes would be sufficient to make sure that the time-scale of the simulations remains the same. To see this, we consider the mean-squared displacement MSD of a monomer in some time $t$:

$$\mathrm{MSD} \propto b^2 \left(\frac{t}{\tau_0}\right)^{1/2}, \tag{S10}$$

where $\tau_0 = b^2 \mathrm{k_B T}/\zeta$ is the time it takes for a monomer of size $b$ to diffuse its own radius. By plugging in the expression for $\tau_0$, we find

$$\mathrm{MSD} \propto b t^{1/2} \left(\frac{\mathrm{k_B T}}{\zeta}\right)^{1/2}. \tag{S11}$$

If two models have the same MSD in times $t$ and $t'$, we have

$$\frac{t}{t'} = \frac{Nb^2}{N'b'^2}. \tag{S12}$$

The RHS is the ratio of the squared end-to-end distances for an ideal chain. Hence, for an ideal chain where we would set $b' = b\sqrt{N/N'}$ to match the dimensions of the polymer, the right hand side would be one. However, for a confined chain with excluded volume interactions, we expect $b' = b(N/N')^{\nu}$ with $\nu \neq 1/2$, and the time-scales of the models would hence not be equal.

In addition to the above changes, to ensure the models have consistent time-scales, we adjust the number of 3D simulation time-steps corresponding to 1 s in the 1D loop-extruder simulations as in previous work [12]. Briefly, we calibrate each model's time-scale by matching experimental data for the short-time mean-squared displacements of labeled loci [18, 19]. After these calibrations, both the short time-scale dynamics and length-scales of our mechanistic model are comparable with each other and in agreement with experimental data (Supplementary Figure S14). The models with different $N$ show similar chromosome organization (Supplementary Figure S15), suggesting that our results are not heavily dependent on the coarse-graining scale. The most notable difference is that the $N = 404$ model shows more global compression of the chromosome (see Local extension in the mechanistic model below).

### 7 Local extension

#### 7.1 Relaxation of relative extension after replication

To assess trends in relative extension over time, we fit exponential curves to the mean relative extensions of different regions after their replication, using least-squares fitting with the Julia package CurveFit (Supplementary Figure S12A). For regions of size $100 - 300$ kb, most data sets are well-fitted with exponentials. All fit time-scales are shown in Table S2.

The mean time-scales of decay for $n = 5, 10, 15$ are $10.2, 9.9, 8.7$ minutes correspondingly. For $n = 20, 25$, fit qualities decreased, and regions near 3250-3400 kb give positive fit exponents. For these region widths, excluding the positive fit exponents, mean time-scales of relaxation are given by 10.5 and 16.0 minutes correspondingly. Using the 4D-MaxEnt method, we hence infer exponentially decaying relative extension on newly replicated chromosomal regions, with decay times of order 10 minutes.

#### 7.2 Relative compression ahead of replication forks

To assess whether and how relative extension changes ahead of the replication forks, we calculate the mean local extension averaged over $50, 100, 150, 200, 250$ kb regions ahead of both forks (Supplementary Figure S12B). As the averaging interval increases, the strain ahead of the replication forks becomes more scattered. For averaging regions of size $50 - 150$ kb, we find that after 10 minutes, the 4D-MaxEnt model shows relative compression ahead of both replication forks. However, no clear trend over time is seen; most data points are scattered close to the mean value.

##### 7.3 Local extension with fewer constraints

To assess what causes the changes seen in the local extension of the full 4D-MaxEnt model, we analyze the local strain patterns of the *ori*-constrained model and the unsegregated model constrained only on Hi-C data.

We find that the *ori*-constrained model shows some extension near the *ori* regions at $t = 10$ min (Supplementary Figure S11A). However, at all other time-points, there is no local strain visible near the replication forks. We hence conclude that *ori*-constraints do not explain the 4D-MaxEnt inference for local extension behind and local compression in front of the replication forks.

The model constrained only using the Hi-C data, on the other hand, displays patterns of local extension around the replication forks (Supplementary Figure S11B). Newly replicated regions show local extension, whereas regions ahead of the replication forks are compressed. Unlike in the full 4D-MaxEnt model, the relative extension near the origin does not seem to decay back to zero over time.

##### 7.4 Local extension in the mechanistic model

To see whether loop extrusion or *ori*-pulling could explain the patterns of local extension inferred using the 4D-MaxEnt approach, we also calculate the relative extensions for the mechanistic models considered in Main Text Figure 3.

We find that all mechanistic models, with or without loop-extruders, show overall compaction relative to $t = 0$ min (Supplementary Figure S11A). This is especially apparent when the number of monomers is small ($N = 404$) (Supplementary Figure S15C). We note that when $N$ is larger, the stiffness of the coarse-grained springs is the lower ($k \sim \mathrm{k_B T}/\sigma^2$, where $\sigma$ is the standard deviation in the spring length).

We first consider how compression arises without loop-extruders. We can attribute the local compression over time to the compression of the coarse-grained springs due to the increasing density of the system. Whereas replication proceeds linearly in time, the cell grows exponentially in time, so that the density of the system follows

$$\rho(t) = \frac{N + 2v_{\mathrm{repl}}t}{V_0 e^{t r_{\mathrm{growth}}}}, \tag{S13}$$

where $V_0$ is the original volume of the system, and $r_g$ is the exponential growth rate of the cell. For $t \leq 75$ min, the density is increasing, and hence the springs in the mechanistic model experience an increasing pressure. As a result, the springs are uniformly compressed, and accordingly, the relative extension averaged over all loci follows a simple scaling law $\langle d_n \rangle_t \,/\, \langle d_n \rangle_{t=0} \propto \rho(t)^{-1/3}$ (Supplementary Figure S15D). For models with a finer coarse-graining level ($N = 1620$ or $N = 4050$), this scaling is still present, although the changes in local compaction are smaller, since the springs are stiffer (Supplementary Figures S15C).

In addition to density-mediated compaction, we find that mechanistic simulations with loop-extruders show more relative compaction than simulations without loop-extruders (Supplementary Figure S11A). This can be explained by changes in the loop size distribution over time. After replication initiation, the density of small loops in the system increases (Supplementary Figure S11B,C). Since small loops cause local compaction, this shift in the loop size distribution results in apparent compaction compared to $t = 0$ min.

We can attribute the decrease in loop sizes during replication to two factors: the binding of new loop-extruders during replication, and increased unbinding and rebinding of loop-extruders due to the movement of the replication forks. We now develop an analytical model that describes the time-dependence of the number of small loops.

Let $n_{\text{small}}$ be the fraction of loop-extruder loops that are below a certain size $l$. The time-evolution of this quantity can be described as

$$\frac{dn_{\text{small}}}{dt} = \frac{1 - n_{\text{small}}}{\tau} + \frac{M_0}{M(t)} \frac{2v_{\text{repl}}}{N} + \frac{D(v_{\text{repl}} t)(1 - n_{\text{small}}) v_{\text{repl}}}{M(t) \Delta x} - n_{\text{small}} k_{\text{grow}}. \tag{S14}$$

The first term on the LHS describes the unbinding and rebinding of loops larger than $l$. The unbinding time $\tau \approx 33$ min is set by the loop-extruder processivity (1200 kb) and leg speed (18 kb/min) [20].

The second term describes the addition of new loop-extruders during replication. Note that this term is decreasing in time; the current number of loop-extruders is given by $M(t) = M_0 + M_0 2 v_{\text{repl}} t / N$, where $v_{\text{repl}}$ is the replication fork speed. Although the absolute rate of loop-extruder addition remains constant ($M_0 2 v_{\text{repl}} / N$), over time these newly added loop-extruders make up a smaller and smaller fraction of the loop-extruder population.

The third term describes unbinding of loop-extruders at the replication forks. Here $D(x)$ is the probability of finding a loop-extruder at lattice site $x$, and $\Delta x$ is the lattice spacing. We multiply the term by the fraction of loops larger than $l$ $(1 - n_{\text{small}})$, since the unbinding of small loops does not change $n_{\text{small}}$. As an approximation, we assume that small and large loops have the same spatial distribution, and that the distribution is symmetric around the origin; $D(x) = D(N - x)$. We also note that there is no factor of two for unbinding at both forks; this is because for each loop-extruder we only consider one leg, since the same loop-extruder cannot unbind twice. The loop-extruder density is highly peaked at the origin (Supplementary Figure S16), meaning that the third rate can vary by orders of magnitude.

Finally, the last term describes growth of loops to sizes beyond $l$. We expect $k_{\text{grow}}$ to be proportional to the loop-extruder stepping speed $v_{\text{LE}}$, but the prefactor will vary with $l$. Furthermore, the expected number of collisions between loop-extruders affects $k_{\text{grow}}$, since loop-extruder stalling upon collision slows down loop growth.

We see that at earlier replication stages, the rate at which $n_{\text{small}}$ increases is larger. This qualitatively matches our findings that there is an initial increase in the fraction of small loops, followed by a decrease. We can also solve (S14) numerically for a given set of parameters. First, we consider simulations without loop-extruder unbinding at the replication forks. In these simulations, the third term in (S14) can be neglected, and the only unknown parameter is $k_{\text{grow}}$. We find that using $k_{\text{grow}} \approx v_{\text{LE}}/\Delta x/17$ gives a good fit to our data for $l = 40$ kb (Supplementary Figure S11D,E). Next, we consider simulations with loop-extruder unbinding at the forks. We use the same value of $k_{\text{grow}} = v_{\text{LE}}/\Delta x/17$. We estimate $D(x)$ with the initial distribution of loop-extruders at the beginning of our simulations. We find that the analytical solution slightly overestimates the number of small loops at early time-points, but otherwise predicts the data well (Supplementary Figure S11C). The over-estimate at early replication stages likely results from the density of small loops being higher near the origin, which results in an over-estimation of replication fork-mediated unbinding of large loops in this region.

We hence conclude that in all our mechanistic MD simulations, the increasing density of the system over time results in global compaction of the chromosome compared to $t = 0$ min. Additionally, in mechanistic models with loop-extruders, replication initially increases the density of small loop-extruder-mediated loops in the system, which further compacts the chromosome relative to $t = 0$ min. Importantly, however, neither loop extrusion or *ori*-pulling gives rise to extension behind and compaction ahead of the replication forks, as inferred using the 4D-MaxEnt method.

*In vivo*, we expect the total compaction level of the chromosome to be affected by mechanisms such as NAP binding, supercoiling, and macromolecular crowding. These factors might all vary over the course of the replication cycle, which is not taken into account in our mechanistic model. Furthermore, our assumption of a linear growth rate for loop-extruder numbers may be incorrect. These considerations could explain why bacterial chromosomes may not show overall compaction during replication *in vivo*, even if the effects seen in our mechanistic model are present.

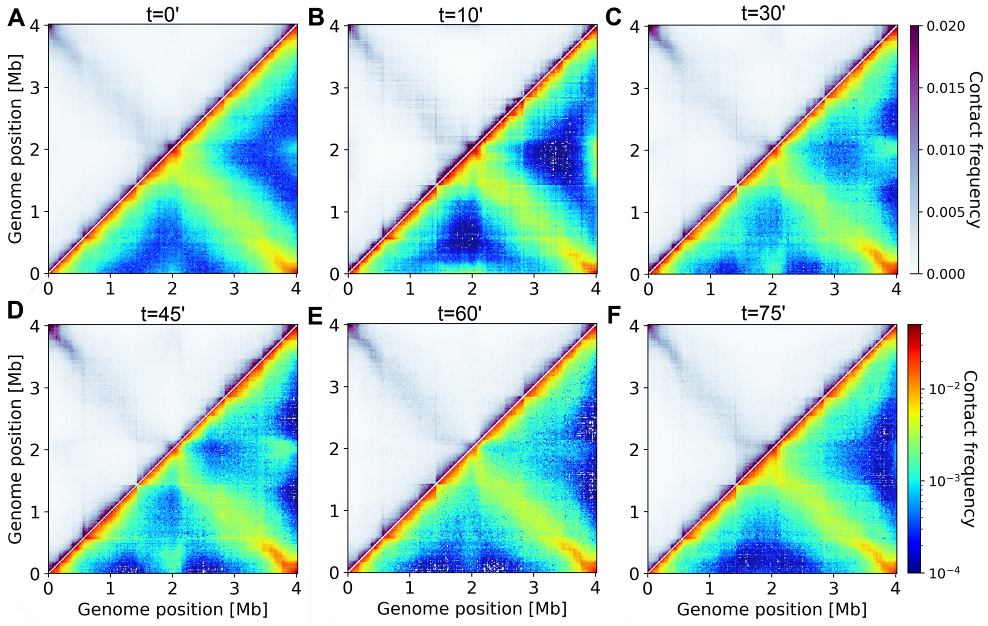

Figure S1: **Input Hi-C maps. A-F**: Input Hi-C maps $f_{ij}^{\text{expt}}(t)$ for *C. crescentus* cells for each cell cycle progression time, obtained via the bias correction procedure applied to data from [5]. Upper left triangles: linear scale. Lower right triangles: logarithmic scale.

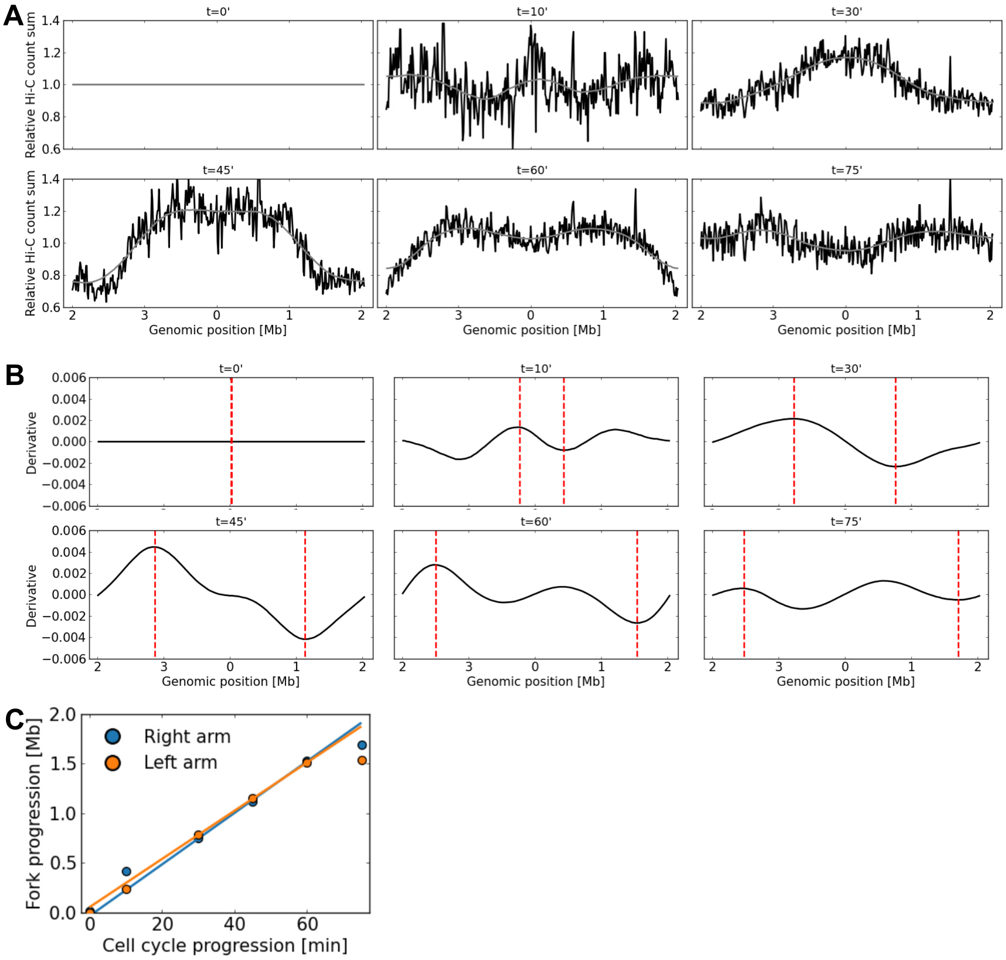

Figure S2: **Inference of fork positions.** (A) Black: column sums $\sum_j f_{ij}^{\text{expt}}(t)$ for each genomic position $i$ and each cell cycle progression time $t$. Gray: smoothened curve, using a Gaussian filter with standard deviation 350 kb. (B) Derivatives of the smoothened curves shown in (A). Dashed lines indicate the estimated replication fork positions for each chromosomal arm, found as maxima and minima of the derivative curves. (C) Linear fits to the inferred fork positions at 30, 45 and 60 minutes.

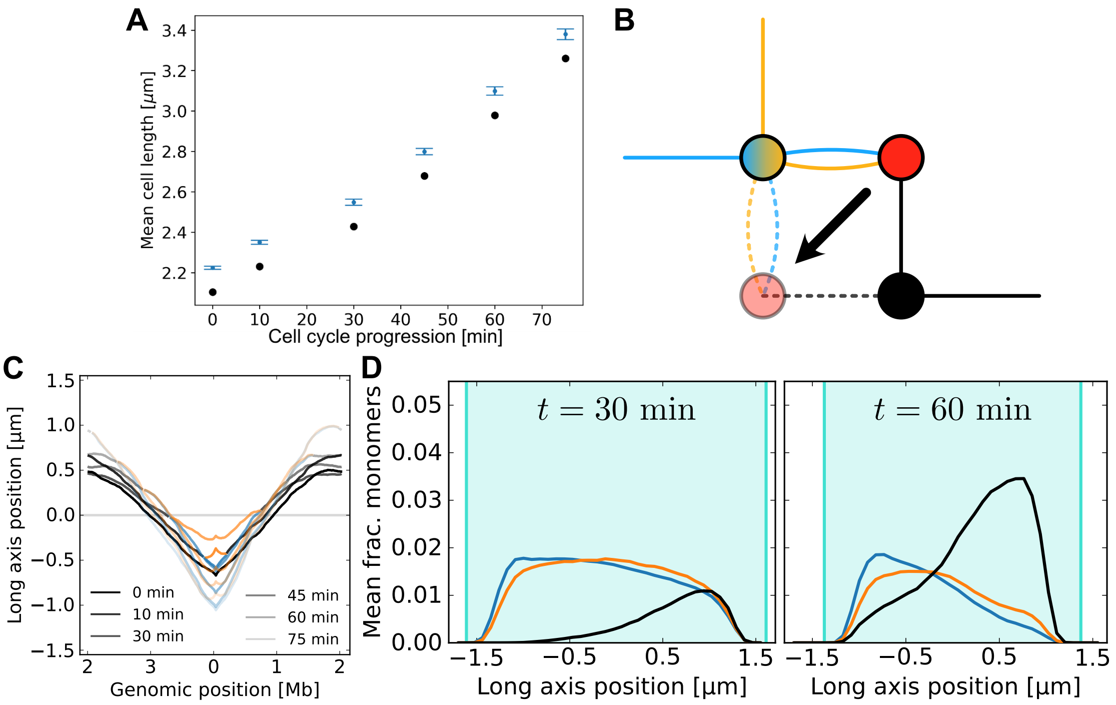

Figure S3: **Construction of model and model without *ori-ori* distance constraints.** (A) Blue dots: mean cell lengths determined for each time since synchronization. The error bars indicate two times the standard error on the mean. Black dots: cell lengths used as 4D-MaxEnt model inputs, with the estimated cell envelope width subtracted from the mean cell lengths. (B) Illustration of the fork move used in the Monte Carlo algorithm. The fork move is performed on the junction site (red) if two of the three connected monomers overlap (orange and blue), and the third monomer (black) is at a 90°angle to the other two. (C) Average long-axis localizations for a model trained only on Hi-C data, similar to Main Text Fig. 3. No segregation is seen. (D) Density histograms of model trained only on Hi-C data, similar to Main Text Fig. 3. The replicated strands colocalize.

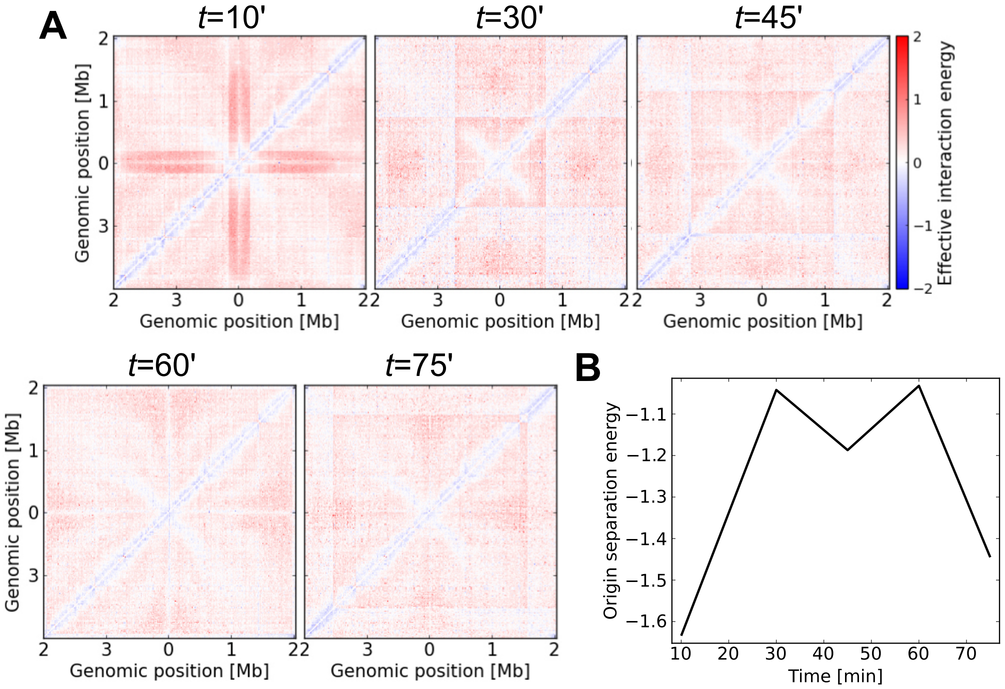

Figure S4: **Effective energies of the 4D-MaxEnt model.** (A) The effective pairwise interaction energies between monomers, $\epsilon_{i,j}$. $t = 0$ min corresponds to the unreplicating model published in [8]. (B) The effective energy $\alpha(t)$ for the *ori*-*ori* separation in the 4D-MaxEnt model.

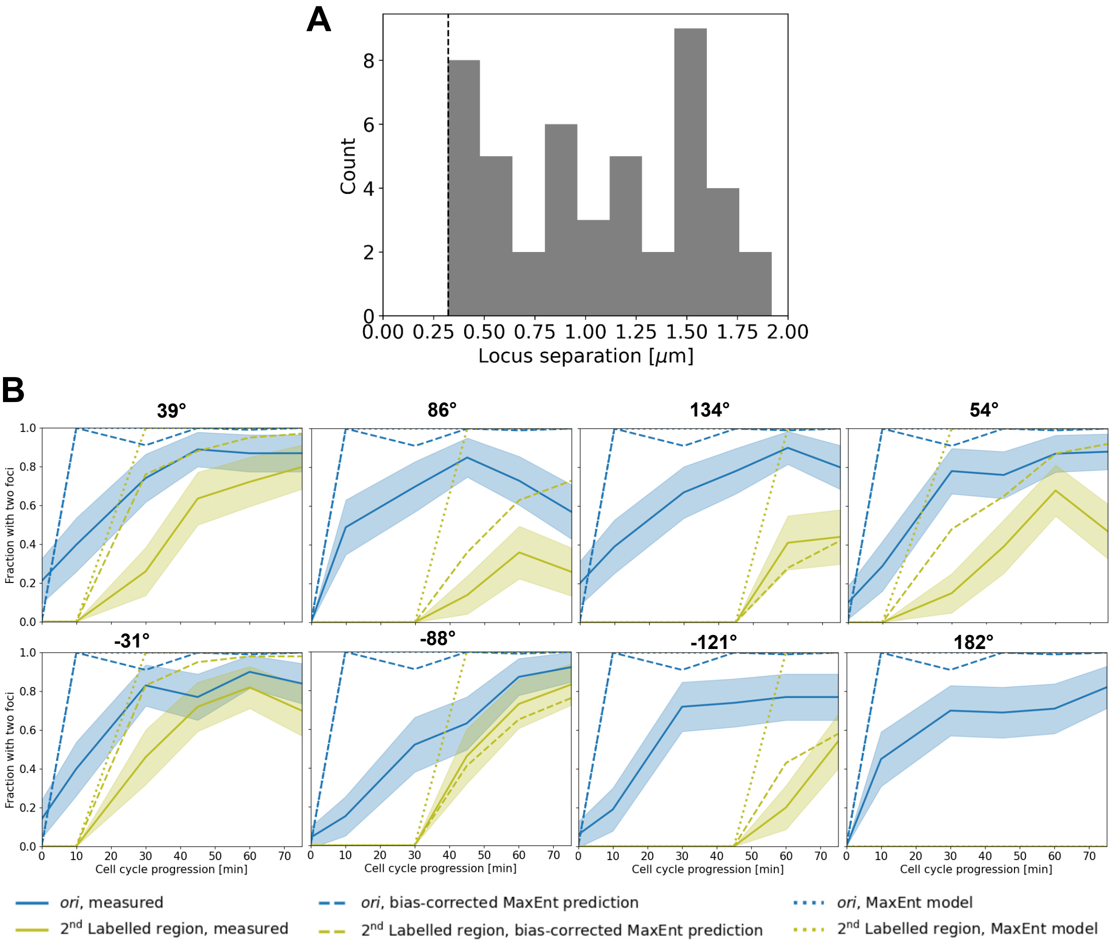

Figure S5: **Effect of resolution limit on measured locus distances.** (A) Histogram of measured focus distances at 86°. Data are taken from all time points from the Tn3 data set. The dashed vertical line indicates the smallest observed distance, equal to 0.32 $\mu$m, which we interpret as the resolution limit. (B) Fraction of cells with two foci for each measured locus, together with value inferred using the 4D-MaxEnt model. Blue solid lines: measured fraction of cells with two *ori*'s. Blue dashed lines: inferred fraction using the 4D-MaxEnt model, given the indistinguishability of two foci at short distance. Blue dotted lines: locus counts in the 4D-MaxEnt model. Yellow solid lines: measured fraction of cells with two copies of the locus indicated at the top left. Yellow dashed lines: inferred fraction for the same region using the 4D-MaxEnt model. Yellow dotted lines: locus counts of the same region in the 4D-MaxEnt model. The shaded areas indicate the error margins (2×SEM) on the experimental data, calculated by assuming binomial sampling.

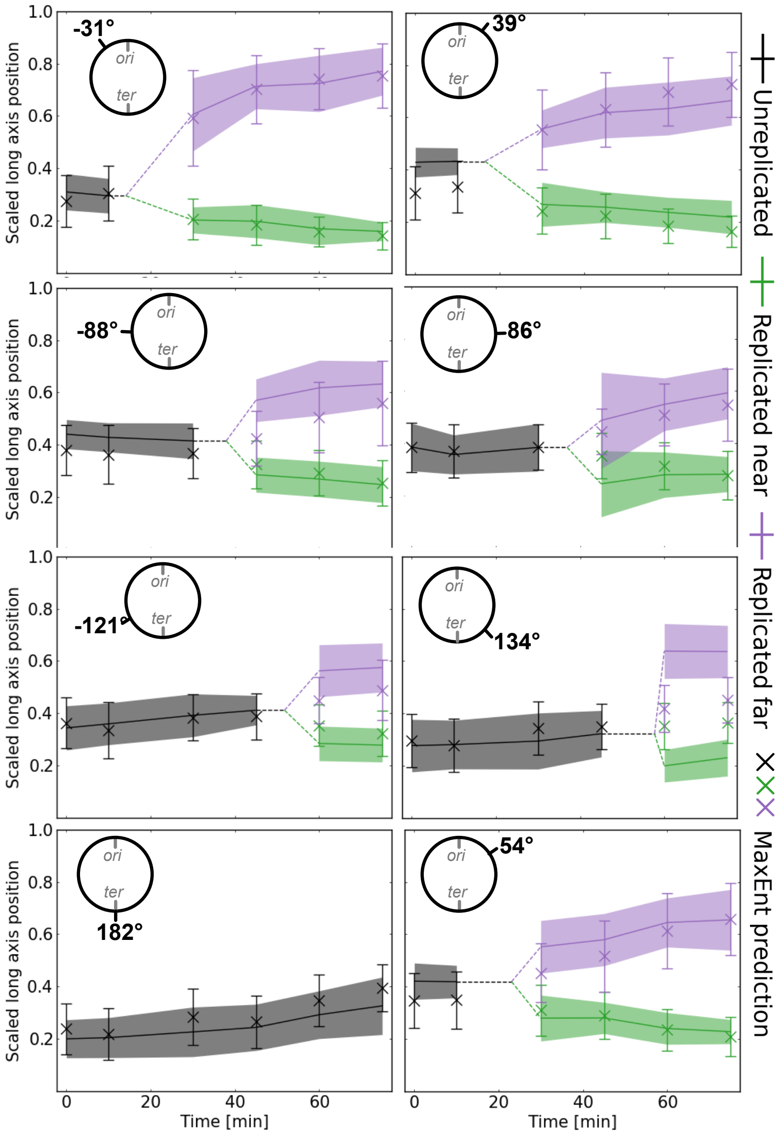

Figure S6: **4D-MaxEnt inferences for validation data without bias-correction.**The inferred distances between replicated foci, similar to Main Text Fig. 2, but without excluding focus pairs at distances below the resolution limit.

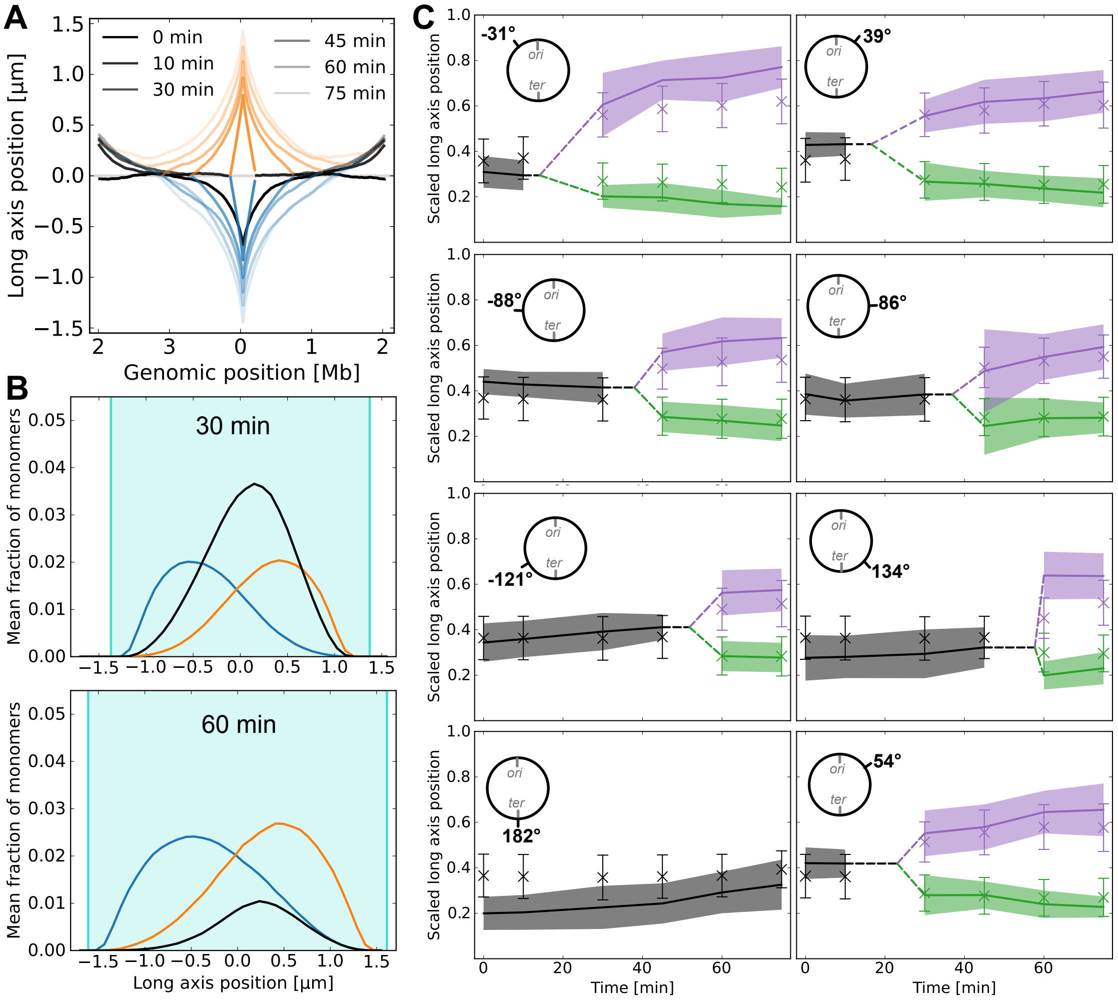

Figure S7: **Origin-constrained model.** (A) The mean long axis positions of loci in the model where only the distance between the origins is constrained. (B) Density histograms at $t = 30$ and 60 min for *ori*-constrained model. (C) *Ori*-constrained model compared to validation data, after bias-correction. The model performs significantly worse than the 4D-MaxEnt model with Hi-C constraints (Main Text Fig.2B).

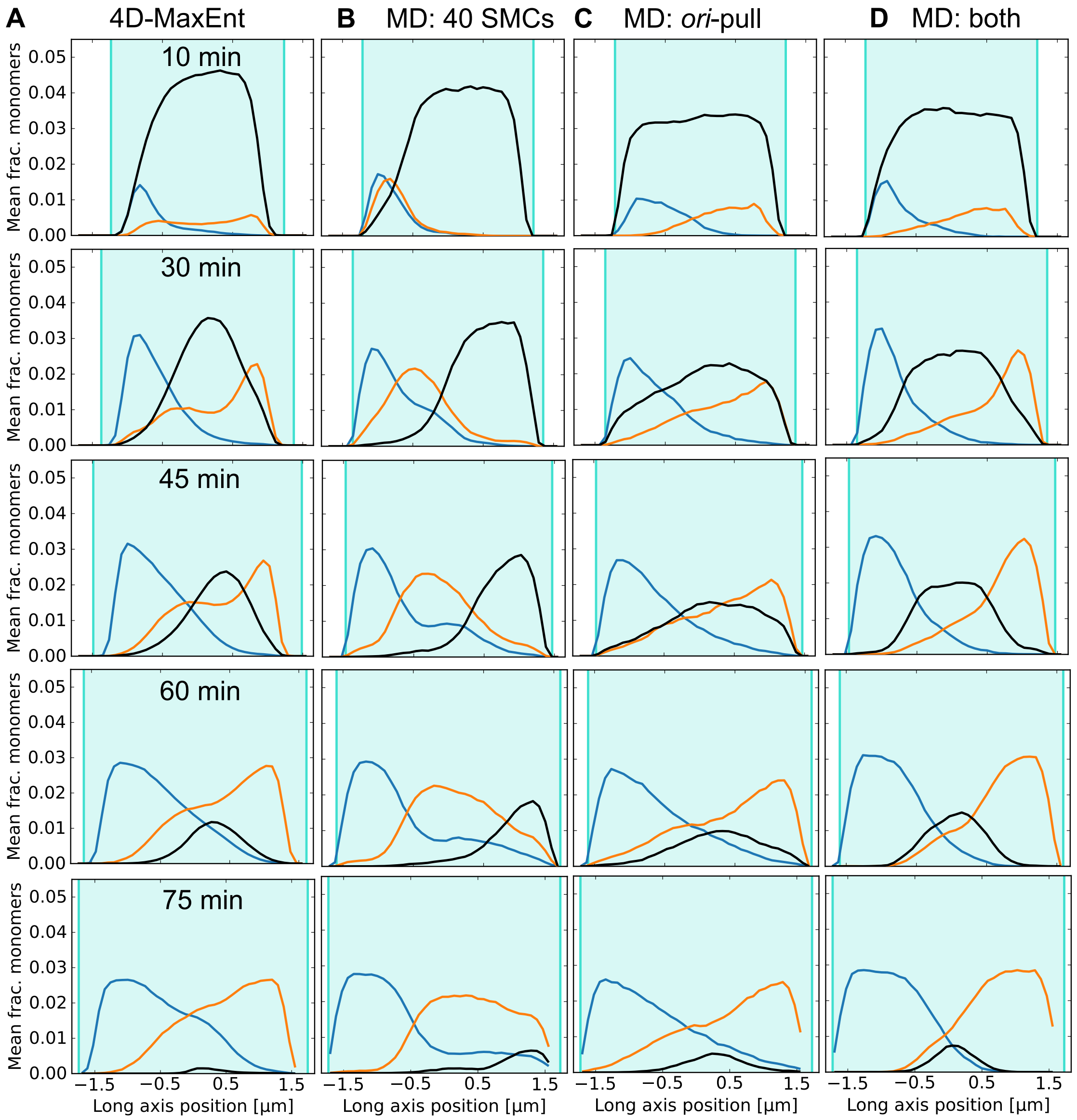

Figure S8: **Density histograms for all time-points.** Density histograms similar to Main Text Fig. 2D-F for all time-points, for the: (A) 4D-MaxEnt model, (B) the mechanistic model with 40 loop-extruders, (C) the mechanistic model with origin-pulling, and (D) the mechanistic model with both origin-pulling and loop-extruders.

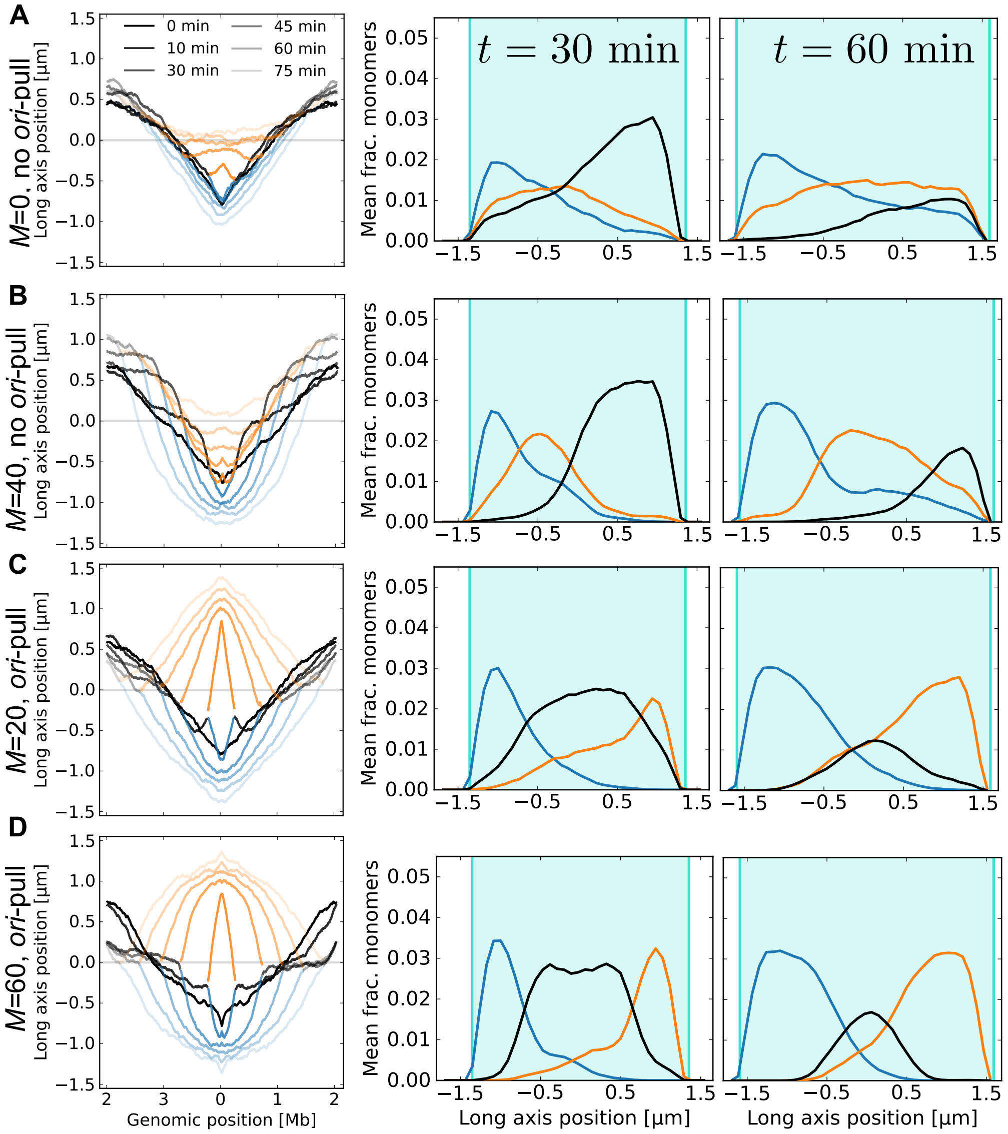

Figure S9: **Fitting the number of loop-extruders per chromosome** $M_0$. Mean long-axis position and density histograms, similar to Main Text Fig. 3, for mechanistic models (A) without loop-extruders or *ori*-pulling, (B) with 40 loop-extruders but no *ori*-pulling, (C) with 20 loop-extruders and *ori*-pulling, and (D) with 60 loop-extruders and *ori*-pulling.

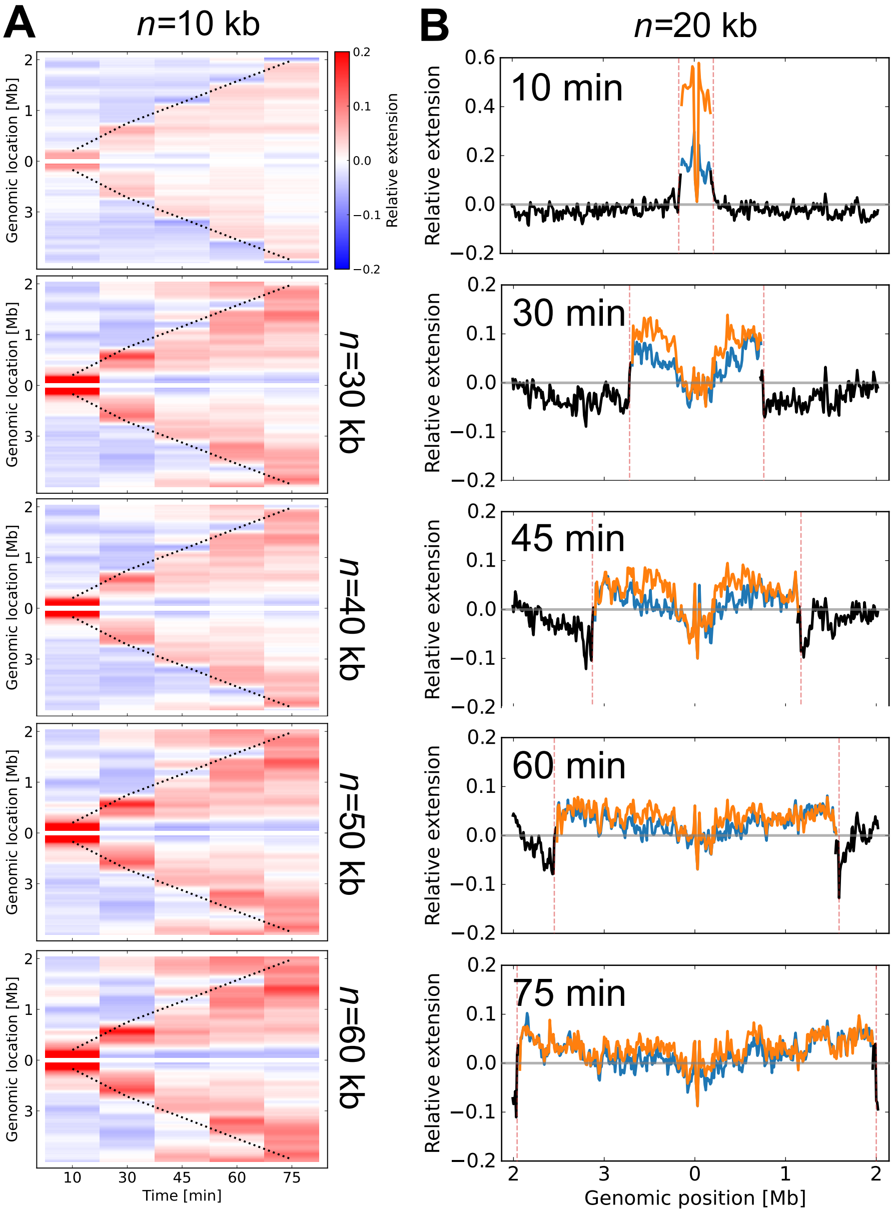

Figure S10: **Relative extension calculated for different** $n$**.** (A) Local extension kymographs similar to Main Fig. 4A, calculated using different values of $n$. Pattern of local extension behind and compaction ahead of the replication forks still visible. (B) The local extension for each time-point, shown for all three strands of the replicating chromosome, using $n = 20$ kb.

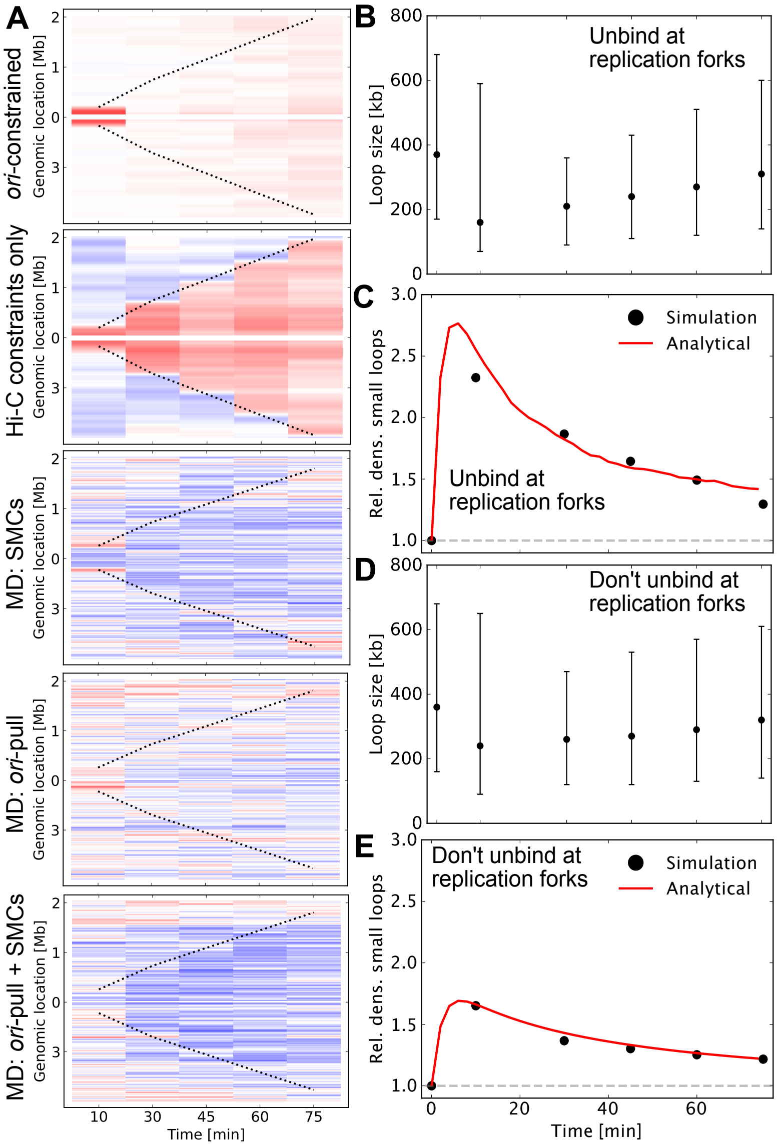

Figure S11: **Relative extensions of other models.** (A) Relative extension kymographs for: data-driven *ori*-constrained model, data-driven model with only Hi-C constraints, mechanistic model with 40 loop-extruders, mechanistic model with *ori*-pulling, and a mechanistic model with both 40 loop-extruders and *ori*-pulling. (B) Median loop size over time in mechanistic simulations with 40 loop-extruders. Error bars indicate 25th and 75th percentile. (C) Density of loops of size $< 40$ kb relative to $t = 0$ min. Black dots: simulations. Red line: numerical solution to Eq. (S14). $k_{\text{grow}}$ was taken from fit in panel E. Loop-extruder distribution was taken from initial density of loop-extruders in simulations. (D) Median loop sizes in mechanistic simulations without loop-extruder unbinding at the replication forks. (E) Relative density of loops of size $< 40$ kb for mechanistic simulations without loop-extruder unbinding at the replication forks. Red line: numerical solution to Eq. (S14), neglecting the third term. The data was used to fit $k_{\text{grow}}$

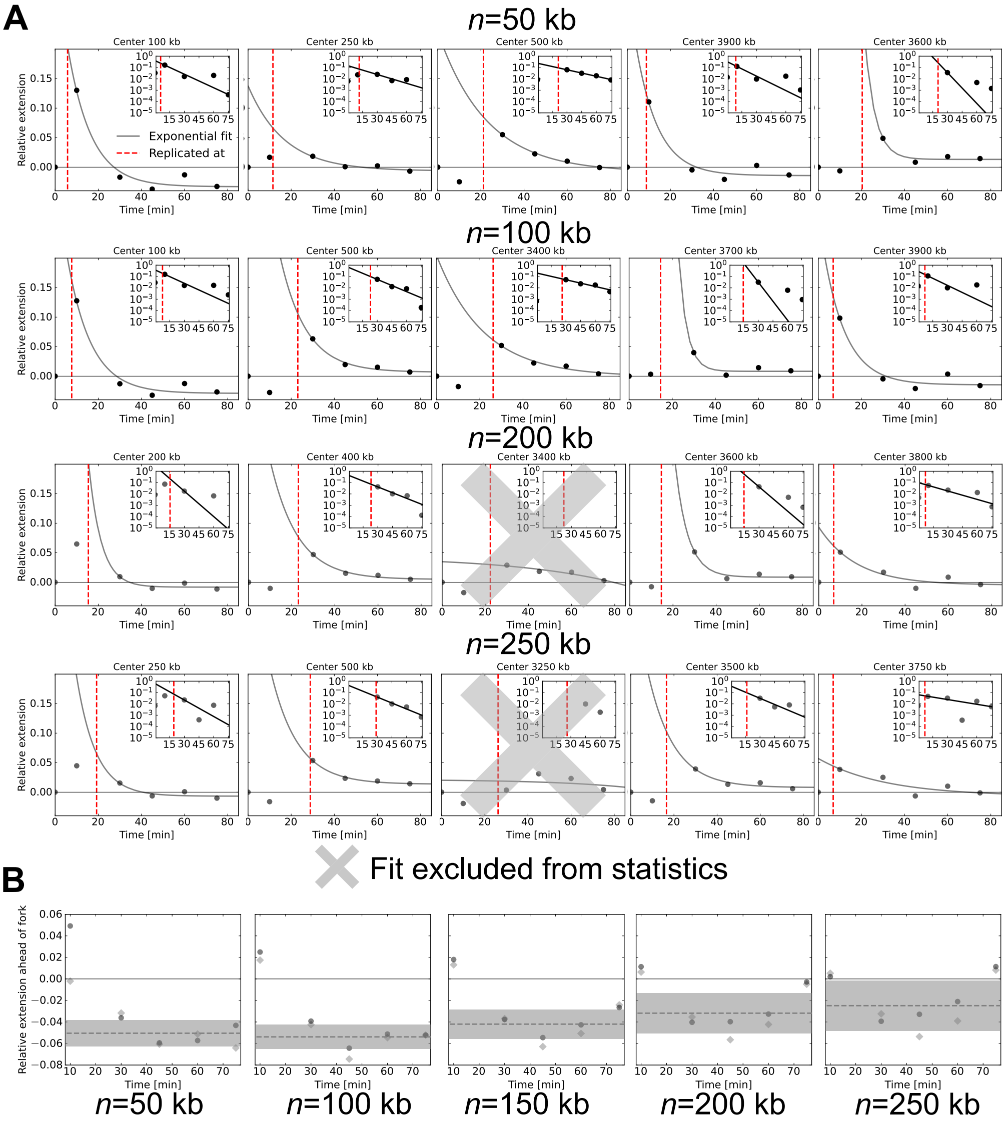

Figure S12: **Analyzing local extension around replication forks.** (A) Example exponential fits to the inferred mean local extension in different regions over time, for varying region width $n$. Fits were performed on with midpoint spacing $n$ with at least four time-points after replication. Red vertical dashed lines indicate when each region was replicated. Horizontal line indicates zeros strain. Top-right insets show the extensions minus fit constant on a logarithmic scale; values below the fit constant not shown. Crossed out plots indicate fits with negative fit time-scales, which were excluded from the mean statistics. For $n \geq 150$kb, negative time-scales were found for the region behind the fork on the left arm of the chromosome. (B) Inferred mean compression ahead of the replication forks. Dashed vertical line indicate mean strain for time-points $t > 10$, error bar indicates standard deviation. For multiple values of $n$, we find no clear trend in the compaction over time.

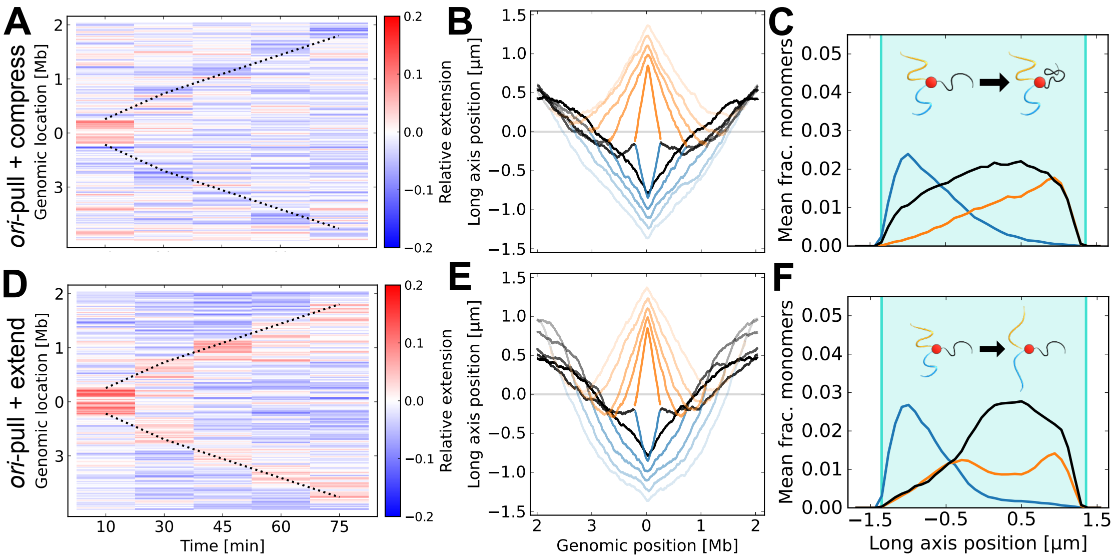

Figure S13: **Mechanistic models with only compression or extension around the forks.** First row: model with *ori*-pulling and only compression ahead of the forks, with panels showing (A) the relative extension kymograph, (B) the mean long-axis position curves, and (C) the density histograms at $t = 30$ min. No extension is seen behind the replication forks. Second row: model with *ori*-pulling and only extension behind the replication forks. Panels show (D) the relative extension kymograph, (E) the mean long-axis position curves, and (F) the density histograms at $t = 30$ min.

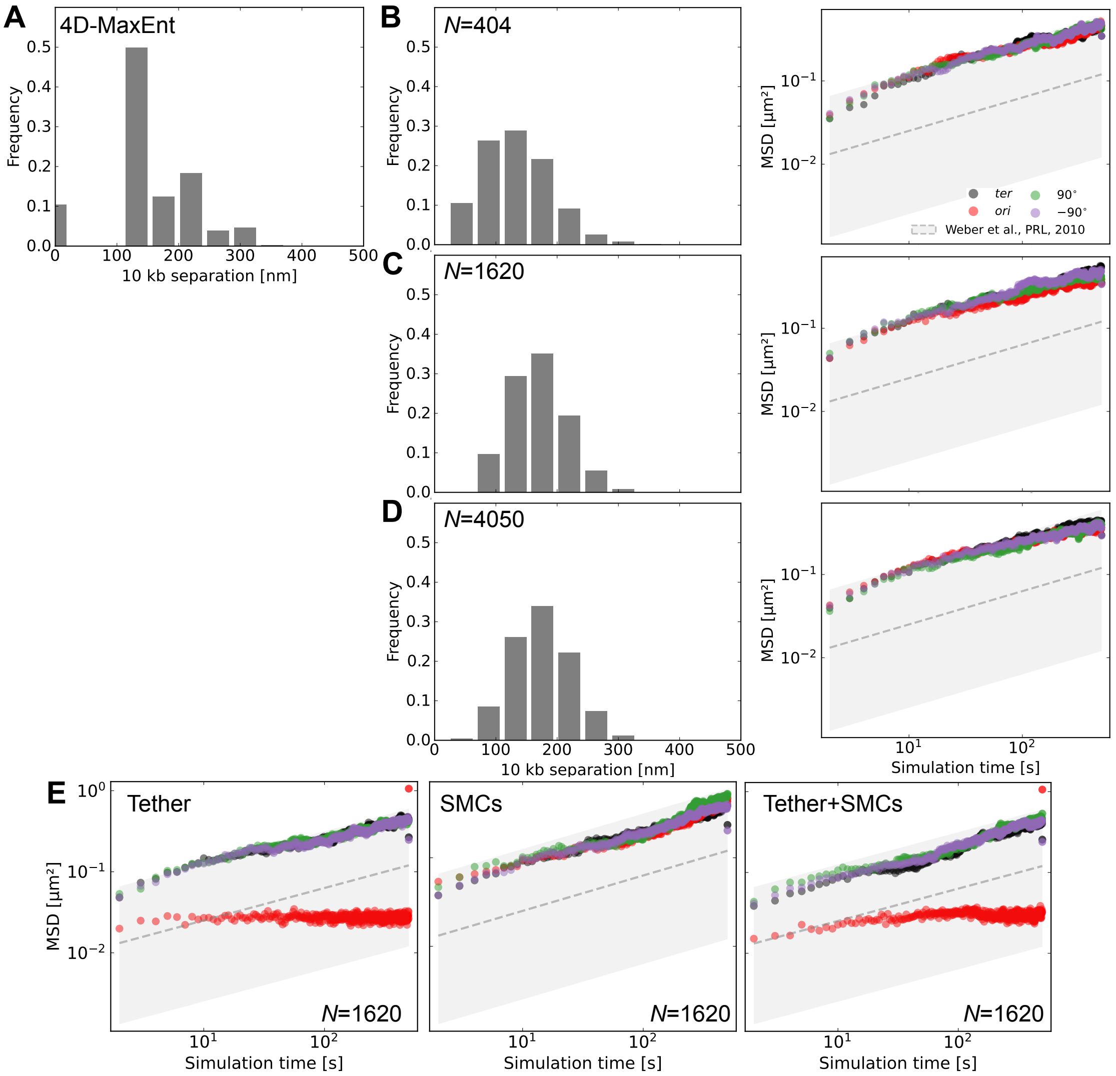

Figure S14: Setting time-scales and length-scales of mechanistic model. (A) Histogram of extensions of 10-kb segments in the 4D-MaxEnt model. (B) For mechanistic model without loop-extruders or *ori*-pulling, using $N = 404$. Left: histogram similar to A. Right: mean-squared displacement (MSD) as a function of time, compared to experimental data [18]. (C) Similarly for mechanistic simulations with $N = 1620$. (D) Similarly for mechanistic simulations with $N = 4050$. (E) MSD as a function of time for simulations with $N = 1620$, with or without *ori*-pulling and/or loop-extruders.

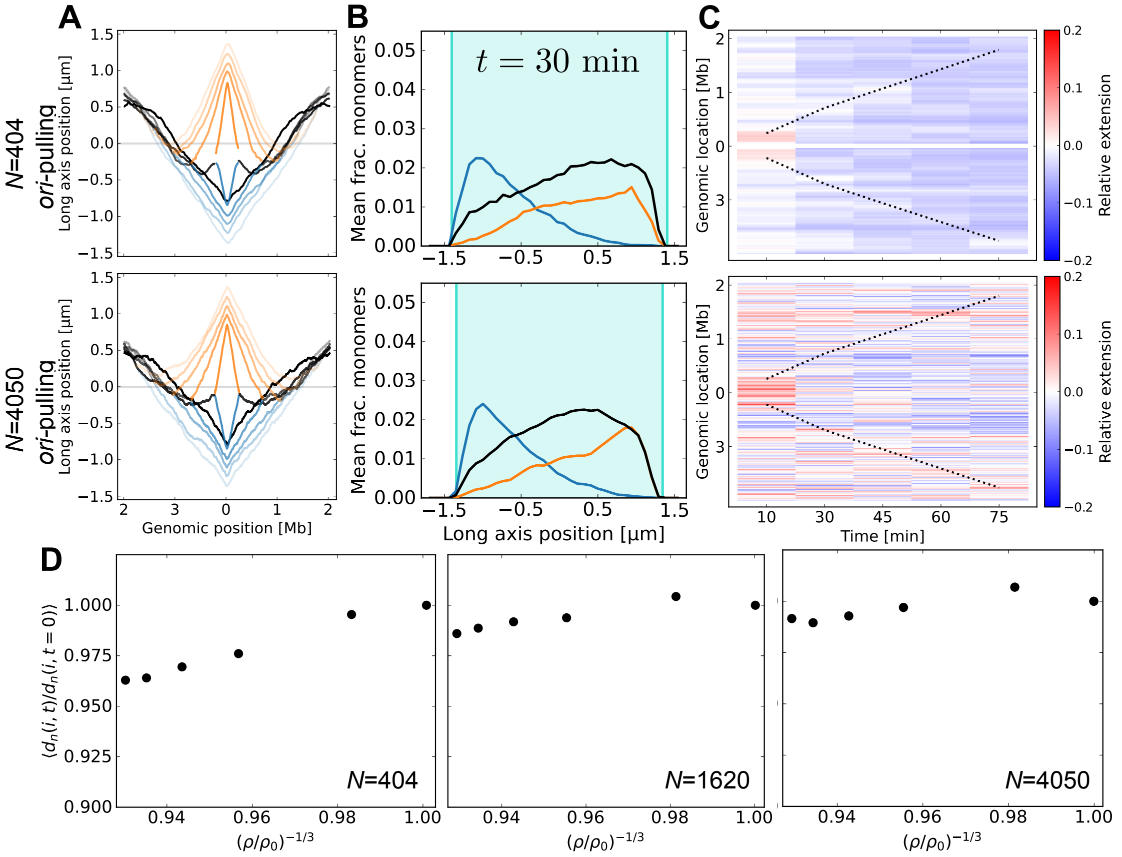

Figure S15: **Mechanistic models with different coarse-graining levels.** (A) Mean long-axis positions of all monomers for mechanistic models with different coarse-graining levels. Comparable to Main Fig. 3B and C. (B) Density histograms at $t = 30$ min for the same models. Comparable to Main Fig. 3E and F. (C) Local extension kymographs for the same models. For $N = 404$, we see an overall compression compared to $t = 0$ min. Comparable to Supplementary Figure S11. (D) Average relative extension $d_n(i,t)/d_n(i,0)$ over the entire chromosome, as a function of the monomer density of the system relative to $t = 0$. Data shown for *ori*-pulling simulations with $N = 404,\ 1620$, or 4050.

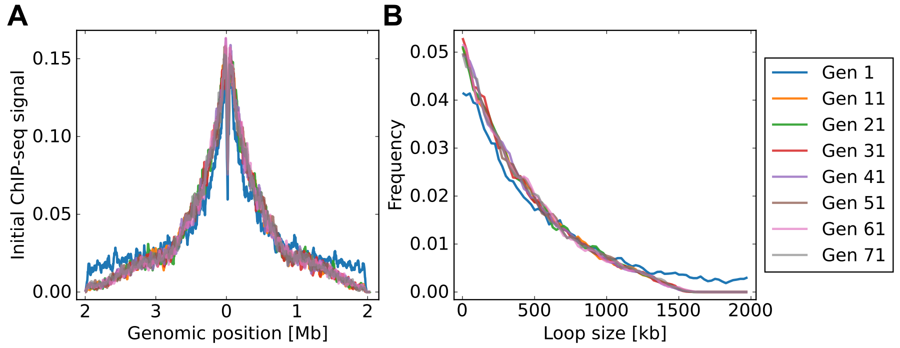

Figure S16: **Loop-extruder distribution inheritance.** In 1D loop-extruder simulations, the loop-extruder positions at $t = 0$ min are inherited from the final positions in the previous simulation. For generation 1, the initial loop-extruder positions are from the steady state distribution without replication. (A) Simulated ChIP-seq signal for loop-extruders at $t = 0$ for different generations. After 10 generations, the distributions look similar. The dip at the *ori* is due to unbinding of loop-extruders at the origin due to replication initiation at $t = 0$. (B) The loop-size distribution at $t = 0$ for different generations. Again, after 10 generations the distribution looks similar. Based on these statistics, we ignore the first 20 generations, so that all simulations have their initial loop-exturder positions sampled from the same distribution.

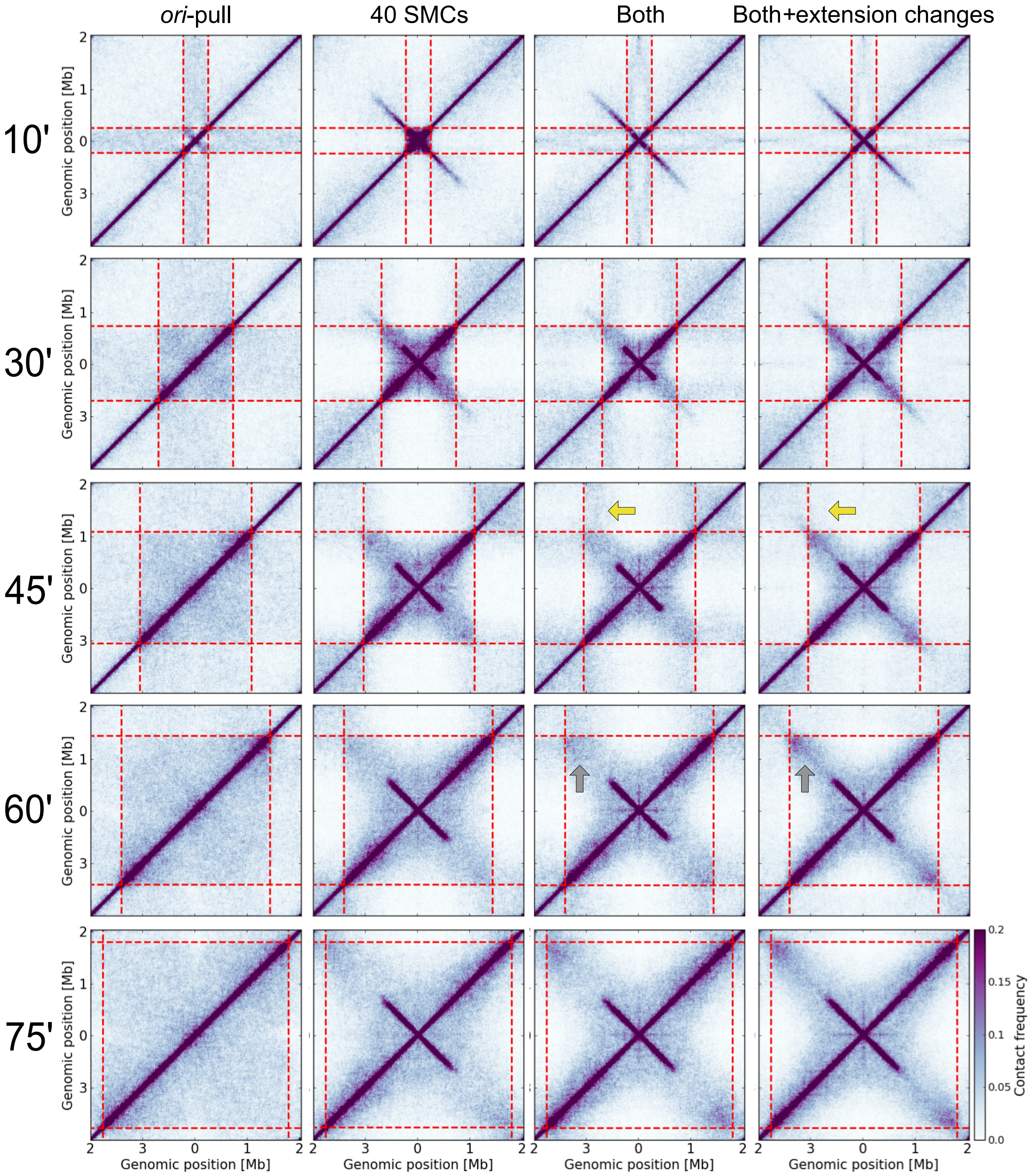

Figure S17: **Contact maps of mechanistic models.** The contact maps (similar to Hi-C) of mechanistic models over time (different rows). Columns correspond to models with only *ori*-pulling, only 40 loop-extruders, both *ori*-pulling and 40 loop-extruders, as well as both *ori*-pulling and 40 loop-extruders, together with extension behind and compaction ahead of the forks. Yellow arrows indicate decrease in contacts between replicated and unreplicated regions after addition of local compaction changes. Gray arrows indicate increase in contacts between newly replicated regions after addition of local compaction changes.

Table S1: *C. crescentus* strains used in this study

| Strain | Genotype/description | Reference |
|---|---|---|
| CB15N | Synchronizable wild-type strain | Evinger & Agabian (1977) [21] |
| MvT171 | CB15N $P_{xyl}$::$P_{xyl}$-*lacI-cfp-tetR-yfp*10x tetO and 10x lacO spaced 10.0 kb apart at 108° | Messelink et al (2021) [8] |
| Tn3 | CB15N $P_{xyl}$::$P_{xyl}$-*lacI-cfp-tetR-yfp* (*lacO*)$_n$ integrated at the *ori* and (*tetO*)$_n$ integrated at bp 957206 (86°) | Viollier et al. (2004) [22] |
| Tn4 | CB15N $P_{xyl}$::$P_{xyl}$-*lacI-cfp-tetR-yfp* (*lacO*)$_n$ integrated at the *ori* and (*tetO*)$_n$ integrated at bp 2026048 (182°) | Viollier et al. (2004) [22] |
| Tn8 | CB15N $P_{xyl}$::$P_{xyl}$-*lacI-cfp-tetR-yfp* (*lacO*)$_n$ integrated at the *ori* and (*tetO*)$_n$ integrated at bp 1498826 (134°) | Viollier et al. (2004) [22] |
| Tn11 | CB15N $P_{xyl}$::$P_{xyl}$-*lacI-cfp-tetR-yfp* (*lacO*)$_n$ integrated at the *ori* and (*tetO*)$_n$ integrated at bp 3029646 (272°) | Viollier et al. (2004) [22] |
| Tn49 | CB15N $P_{xyl}$::$P_{xyl}$-*lacI-cfp-tetR-yfp* (*lacO*)$_n$ integrated at the *ori* and (*tetO*)$_n$ integrated at bp 596575 (54°) | Viollier et al. (2004) [22] |
| Tn72 | CB15N $P_{xyl}$::$P_{xyl}$-*lacI-cfp-tetR-yfp* (*lacO*)$_n$ integrated at the *ori* and (*tetO*)$_n$ integrated at bp 2673431 (239°) | Viollier et al. (2004) [22] |
| Tn85 | CB15N $P_{xyl}$::$P_{xyl}$-*lacI-cfp-tetR-yfp* (*lacO*)$_n$ integrated at the *ori* and (*tetO*)$_n$ integrated at bp 433392 (39°) | Viollier et al. (2004) [22] |
| Tn102 | CB15N $P_{xyl}$::$P_{xyl}$-*lacI-cfp-tetR-yfp* (*lacO*)$_n$ integrated at the *ori* and (*tetO*)$_n$ integrated at bp 3645906 (329°) | Viollier et al. (2004) [22] |

| **Center position** $i$ **[kb]** | **Region width** $n$ **[kb]** | **Fit timescale [min]** |
|---|---|---|
| 50 | 50 | 10.978369538060399 |
| 100 | 50 | 10.893278767354523 |
| 150 | 50 | 10.945396955138651 |
| 200 | 50 | 13.853393191849307 |
| 250 | 50 | 16.796574350009333 |
| 300 | 50 | 13.455461929022821 |
| 350 | 50 | 14.396917500091199 |
| 400 | 50 | 11.078680459628568 |
| 450 | 50 | 17.327448831259883 |
| 500 | 50 | 22.69307843883503 |
| 550 | 50 | 10.876761057695397 |
| 600 | 50 | 7.285833714950176 |
| 650 | 50 | 7.831256057301041 |
| 700 | 50 | 20.59169588912098 |
| 3400 | 50 | 10.194523875783602 |
| 3450 | 50 | 8.438941719367877 |
| 3500 | 50 | 8.918320485365355 |
| 3550 | 50 | 6.570817964724809 |
| 3600 | 50 | 4.795618054188173 |
| 3650 | 50 | 2.8922877045592443 |
| 3700 | 50 | 3.9253001936117506 |
| 3750 | 50 | 6.233170176060974 |
| 3800 | 50 | 4.717984258136371 |
| 3850 | 50 | 1.5516363342746546 |
| 3900 | 50 | 10.227515426403402 |
| 3950 | 50 | 9.164449200865011 |
| 4000 | 50 | 9.334942763636153 |
| 100 | 100 | 11.013338066217326 |
| 200 | 100 | 2.3365556626638835 |
| 300 | 100 | 14.24029964746172 |
| 400 | 100 | 15.96320404692256 |
| 500 | 100 | 12.034977436196284 |
| 600 | 100 | 10.177721019512829 |
| 3400 | 100 | 21.887692327641247 |
| 3500 | 100 | 7.526451100494726 |
| 3600 | 100 | 4.3984023729361175 |
| 3700 | 100 | 3.8747613130973586 |
| 3800 | 100 | 4.608152589175087 |
| 3900 | 100 | 10.486589415883271 |
| 150 | 150 | 1.5436264769161299 |
| 300 | 150 | 8.67610207541583 |
| 450 | 150 | 12.557441227977986 |
| 600 | 150 | 12.658994197385505 |
| 3300 | 150 | -14.010441323579837 |
| 3450 | 150 | 13.211822769697044 |
| 3600 | 150 | 6.2215256717584575 |
| 3750 | 150 | 3.2821742800416525 |
| 3900 | 150 | 11.201021267339593 |
| 200 | 200 | 5.820751954452474 |
| 400 | 200 | 12.691492993286811 |
| 3400 | 200 | -48.91032213939398 |
| 3600 | 200 | 5.913303310587803 |
| 3800 | 200 | 17.615446793492445 |
| 250 | 250 | 8.975993427140072 |
| 500 | 250 | 12.362159643827463 |
| 3250 | 250 | -35.88648941525042 |
| 3500 | 250 | 11.958500324050236 |
| 3750 | 250 | 30.88104249308266 |

Table S2: Fitted timescales of extension relaxation after replication.
